# Spatial-scERA: A method for reconstructing spatial single-cell enhancer activity in multicellular organisms

**DOI:** 10.1101/2024.10.02.616294

**Authors:** Baptiste Alberti, Séverine Vincent, Isabelle Stévant, Damien Lajoignie, Hélène Tarayre, Paul Villoutreix, Yad Ghavi-Helm

## Abstract

Enhancers play an essential role in developmental processes by orchestrating the spatial and temporal regulation of gene expression. However, mapping the location of these regulatory elements in the genome and precisely characterizing their spatial and temporal activity remain important challenges. Here we introduce a novel *in vivo* and *in silico* method for spatial single-cell enhancer-reporter assays (spatial-scERA) designed to reconstruct the spatial activity of multiple candidate enhancer regions in parallel in a multicellular organism. Spatial-scERA integrates massively parallel reporter assays coupled with single-cell RNA sequencing (scRNA-seq) and spatial reconstruction using optimal transport, to map cell-type-specific enhancer activity at the single-cell level on a 3D virtual representation of the sample. We evaluated spatial-scERA in stage 6 *Drosophila* embryos using 25 candidate enhancers (including 19 uncharacterized regions), and validated the robustness of our predicted reconstructions by comparing them to microscopy images generated by *in situ* hybridization. Remarkably, spatial-scERA faithfully reconstructed the spatial activity of these enhancers, even when the enhancer-reporter construct was expressed in as few as 10 cells. Our results demonstrate the importance of integrating transcriptomic and spatial data for the accurate prediction of enhancer activity patterns in complex multicellular samples. Indeed, we found that chromatin modifications and open chromatin regions are often poor predictors of enhancer activity. Moreover, spatial data can often be essential for the accurate annotation of scRNA-seq clusters. Finally, we showed that spatial-scERA could be a powerful tool to link enhancers with their potential target genes. Overall, spatial-scERA provides a scalable approach to map spatio-temporal enhancer activity at single-cell resolution without the need for imaging or *a priori* knowledge of embryology and can be applied to any multicellular organism amenable to transgenesis.

## INTRODUCTION

In metazoans, development is a highly regulated process that transforms a single cell to a fully-formed adult organism. This transformation occurs through differentiation processes largely driven by the regulation of gene expression. A key mechanism in this regulation involves non-coding DNA elements known as *cis*-regulatory modules (CRM) such as enhancers, which confer to genes their specific spatial and temporal patterns of expression (1–4). A major challenge in the field remains the comprehensive identification of enhancer sequences and the detailed characterization of their spatio-temporal activity. Although a growing number of enhancers have been functionally characterized in model organisms (5–9), typically through enhancer-reporter assays (3,10,11), the sheer number of putative developmental enhancers far exceeds the number of genes expressed during development. As a result, we are still far from having a complete understanding of the mechanisms by which enhancers regulate gene expression.

Thanks to substantial efforts involving the mapping of DNase I hypersensitive sites, histone modifications, and transcription factor binding sites, the identification of putative enhancers is being discovered at a rapid pace (12–15). The advent of massively parallel reporter assays (MPRA) has further accelerated this process, enabling the testing of thousands of such enhancer candidates in cell culture systems (16– 18). However, these approaches are limited in their ability to provide information about the actual spatio-temporal activity of enhancers in multicellular organisms. Traditional *in vivo* reporter assays, while capable of systematically testing enhancer activity, have low throughput, as each candidate must be tested individually. Although recent attempts to scale up these assays are promising, they are often limited to a few cell types (19), rely on episomal vectors that lack genomic context (20), or require specialized equipment and extensive prior knowledge of the species’ embryology (5,21).

The advent of single-cell sequencing technologies has paved the way for high-throughput enhancer identification at the cellular level in multicellular organisms. For example, recent profiling of single-cell chromatin accessibility during *Drosophila* embryogenesis has revealed over 100,000 open chromatin regions with potential enhancer function at specific developmental stages (22). However, chromatin accessibility remains only a predictive tool for enhancer function and does not always correlate with spatio-temporal activity (23). By combining the efficiency of classical reporter assays with the cellular resolution of single-cell RNA sequencing (scRNA-seq), it is now possible to enhance these predictions, addressing both the throughput limitations of traditional reporter assays and the lack of *in vivo* insights provided by MPRAs. Nevertheless, single-cell reporter assays have so far been limited to cell culture systems (24,25), hence still lacking spatio-temporal resolution. Ultimately, none of these methods offers an unbiased, scalable means of characterizing spatiotemporal enhancer activity at single-cell resolution.

Here, we addressed this limitation by developing spatial-scERA, an *in vivo* single-cell reporter assay that predicts the tissue-specific activity of candidate enhancer regions in a multicellular organism. Our key innovation is the combination of scRNA-seq with spatialization based on optimal transport to reconstruct the activity of enhancers in a virtual embryo without the need for imaging (26,27). We applied this method to 25 candidate enhancers in stage 6 *Drosophila* embryos and demonstrated the robustness of our method by comparing our reconstructions with traditional enhancer-reporter assays coupled with imaging. Our results highlight the importance of spatial data integration to reconstruct the activity of enhancers in complex tissues and predict the enhancer’s target genes.

## RESULTS

### Spatial-scERA reconstructs *in vivo* single-cell enhancer activity in multicellular organisms

To gain a comprehensive view of spatial enhancer activity at single-cell resolution in multicellular organisms, we developed a method that combines MPRAs, single-cell transcriptomics and spatial reconstruction using optimal transport (Fig. 1A-C). This integrated approach, termed spatial-scERA (spatial single-cell Enhancer-Reporter Assay sequencing), covers all the steps from selection of putative enhancers to spatial reconstruction of their activity using computational approaches.

**Fig. 1.**
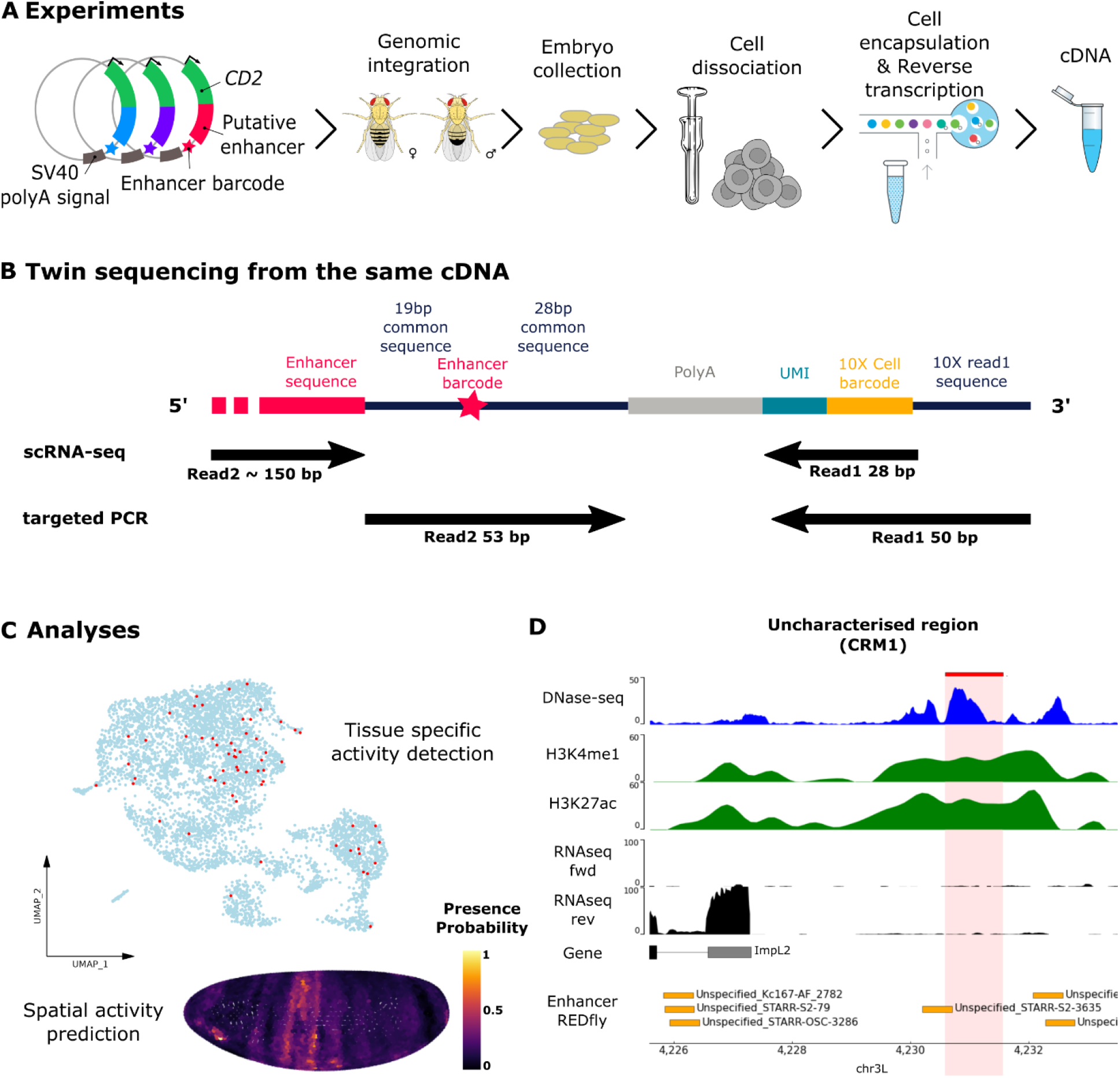
Overview of Spatial-scERA to characterize spatial enhancer activity in *Drosophil*a embryos. (A-C) Overall design of spatial-scERA. (A) A library of enhancer-reporter assay vectors is injected as a pool in *Drosophila* embryos. Genomic integration of the reporter construct is established by screening for red-eyed flies. Embryos are collected from transgenic flies, their cells dissociated and subjected to 3’end scRNA-seq. (B) The cDNA generated during scRNA-seq library preparation is used to generate 2 different sequencing libraries: classical scRNA-seq and targeted PCR amplification. Arrows indicate the portion of the transcript sequenced in each strategy. The location of the enhancer-specific barcode is indicated by a star. (C) Bioinformatic analysis of the sequencing data is used to identify the cell-types in which each enhancer is active, and to generate a spatial reconstruction of each enhancers’ activity in a virtual *Drosophila* embryo. (D) Genome browser plot for one candidate region (highlighted in red). Top to bottom: DNase-seq signal at stage 5 (blue) (15); ChIP-seq signal for H3K4me1 and H3K27ac histone modifications (0-4 hours after egg lay; green) (29,30), RNA-seq signal (2-4 hours after egg lay; black) (31). The location of nearby genes and characterized enhancers from the REDfly database (orange) (32) are indicated.

Spatial-scERA is based on the construction of a reporter library, where each candidate enhancer drives the expression of a reporter gene. As in STARR-seq (18), each candidate enhancer is cloned downstream of the reporter and upstream of a polyadenylation signal, placing the candidate enhancer sequence within the 3’ UTR of the reporter transcript. However, unlike MPRAs, the reporter library is not transfected into cells in culture but directly injected into *Drosophila melanogaster* embryos and integrated in the genome at a single location. As a consequence, screening the expression of the reporter gene provides not only information on the ability of a candidate enhancer to drive expression, but also give key insights regarding its tissue-specific activity within a complex multicellular organism. For the method to be scalable, cell-type-specific expression of the reporter is not established by microscopy as in traditional enhancer-reporter assays, but by detecting the sequence of each candidate enhancer within the reporter transcript in a 3’ scRNA-seq experiment. A key innovation of our method lies in the addition of a specific barcode at the 3’ end of the enhancer sequence. This barcode is used to identify the cells in which a candidate enhancer is active by targeted PCR, significantly increasing the number of positive cells detected in a single experiment. Finally, the *in vivo* spatial activity of each enhancer is predicted by reconstructing their activity pattern on a 3D virtual embryo using a custom version of novoSpaRc, a spatialization method based on optimal transport (27). This customization allows us to combine information from scRNA-seq and targeted PCR amplification, but also handle the sparce nature of the data by mapping the probability of presence of the enhancer at any position within the embryo.

### Selecting candidate enhancers

We evaluated spatial-scERA in stage 6 *Drosophila* embryos using a set of candidate enhancers including both positive controls known to be active at stage 6 and uncharacterized regions. To enrich this pool for potential tissue-specific enhancers active at stage 6, we selected the candidate enhancers based on three parameters (see Methods): (1) their proximity to genes expressed in a tissue-specific manner in stage 6 *Drosophila* embryos (28), (2) their overlap with DNase I hypersensitive regions, and (3) the presence of histone modifications characteristic of enhancer regions (H3K27ac and H3K4me1) (29,30) (Fig. 1D). Importantly, we excluded regions that appeared to be actively transcribed in a bulk RNA-seq dataset (31) to ensure that sequencing reads mapping to the enhancer sequence were specific to the reporter construct. These criteria yielded a total of 111 regions, from which we further selected 25 candidates (Supplementary Table 1) by cross-referencing the candidate regions with a database of characterized *Drosophila* enhancers (32) to include a majority of uncharacterized regions. Our final list included five well-characterized stage 6 enhancers such as the twi_ChIP-42 enhancer regulating the *twist* (*twi*) gene (33,34) and the h_stripe1 enhancer regulating the *hairy* (*h*) gene (35), which were used as positive controls. One enhancer named ChIP-27 was selected as negative control with no activity established at stage 6 (33). The remaining 19 regions were composed of 6 enhancers with established activity at other stages of development but with no information at stage 6, such as the prd01 enhancer active in the posterior half of each segment of the embryo at stage 10 (36). There was no information available regarding the spatio-temporal activity of the remaining 13 regions, which were named CRM1 to CRM13 (Supplementary Table 1).

### Applying spatial-scERA to 25 candidate enhancers

For each of the 25 candidate enhancers, we narrowed down a minimal region of about 1 kb centered around the DNase I hypersensitivity signal (Supplementary Fig. S1-S13). These regions were amplified by PCR, flanking their 3’ end by a 19 bp common sequence (for downstream targeted PCR amplification) and a unique 6 bp enhancer barcode. This cassette was then cloned in a reporter vector downstream of the *CD2* reporter gene under the control of the minimal *hsp70* promoter and upstream of the *SV40* polyadenylation signal (Supplementary Fig. S14). The reporter vector also contains an *attB* site, enabling precise site-specific integration of the vector in the genome of *attP*-containing fly lines using the PhiC31 integrase system. Compared to random integration, targeted integration at the same genomic location prevents artifacts resulting from positional effects. Moreover, the integration process is irreversible and excludes the possibility of multiple integration events. Finally, the reporter vector contains the *mini-white* cassette, allowing for the convenient identification of transgenic flies by the presence of red eyes in the progeny of the injected flies.

We injected a reporter plasmid library containing the 25 candidate enhancers in about 3,250 embryos and crossed the resulting flies in batch to a white-eyed fly line. In the next generation, we obtained 285 red-eyed transgenic animals, which were further crossed to each other to generate a pool of flies containing 2 copies of the reporter construct, one on each allele. All cloning, injection, and fly crossing steps were carried out in batch to allow for the rapid generation of spatial-scERA libraries from multiple candidate enhancers in parallel. We then collected embryos at 2.5 to 3.5 hours after egg lay (corresponding to developmental stages 5 to 7, with a majority of stage 6) from our pool of transgenic flies, dissociated them to a single cell suspension, and profiled gene expression using 3’ end droplet-based scRNA-seq. To increase the number of analyzed cells, we collected and sequenced 3 different batches of embryos from the same pool of transgenic flies.

Following sequencing, we mapped the reads to a modified version of the *Drosophila* genome that included the sequence of our reporter constructs. This resulted in a total of 6,588 high-quality cells from the 3 batches of sequencing, which were grouped into 16 distinct clusters by unsupervised clustering. We labeled the clusters based on the most frequent cell-type annotation in the Berkeley *Drosophila* Genome Project (BDGP) database (37) of the top 10 differentially expressed genes for each cluster. This process resulted in the fusion of these clusters into 11 different cell types, which encompass both major cell-types such as the ectoderm and the mesoderm as well as rarer cell-types such as pole cells and the mesectoderm (Fig. 2A, B). For example, the yolk was characterized by the expression of the gene *sisterless A* (*sisA*) (38) while the *Ptx1* and *stumps* genes were more highly expressed in the endoderm and mesoderm clusters respectively (39,40) (Fig. 2C, Supplementary Fig. S15).

**Fig. 2.**
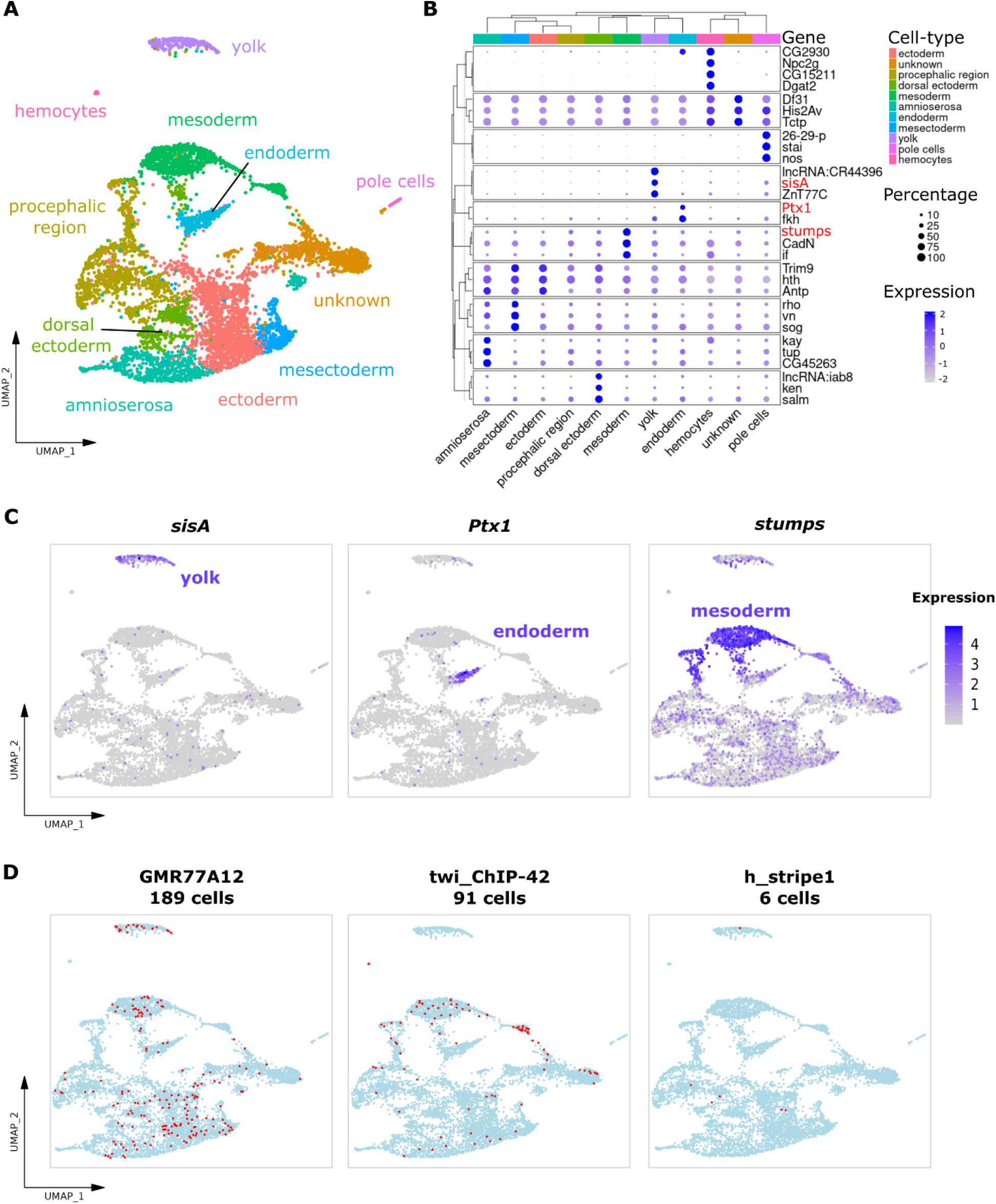
Spatial-scERA can detect tissue-specific enhancer activity. (**A**) Uniform Manifold Approximation and Projection (UMAP) of all cells (n=6,588 cells) present in the scRNA-seq dataset with the 11 identified cell-types. (**B**) Clustered Dot Plot presenting the most differentially expressed genes of each cluster. The size of the dot represents the percentage of cells expressing the gene in each cluster and the color the average differential expression of this gene in this cluster (log2 fold change). (**C**) Examples of highly variable genes selected from the Clustered Dot Plot. The color of the dot represents the expression of a given gene (log2 fold change). (**D**) UMAPs representing the activity of three enhancers. Cells with an active enhancer are shown in red.

### Enhancing recovery through targeted PCR amplification

Having generated a high-quality scRNA-seq dataset, we proceeded to identify the cells expressing our candidate enhancers. In scRNA-seq experiments, detecting lowly expressed transcripts is particularly challenging due to the limited amount of RNA per cell but also because of technical issues such as dropout effects (41,42). As a result, capturing weakly active enhancers can be particularly difficult. To maximize the detection of reporter expression across more cells, we developed a complementary approach that involves targeted PCR amplification and sequencing of a unique enhancer barcode within the reporter construct. This barcode allowed us to confirm and identify new cell-enhancer pairs within the cells sequenced in the scRNA-seq dataset. The 6 bp-long enhancer barcode is located downstream of the enhancer sequence, just upstream of the polyadenylation signal, and is flanked on each side by 2 common sequences absent in the *Drosophila* genome. By using the first common sequence and the 10X Read1 sequence as PCR primers, we generated a 120-bp product comprising both the enhancer barcode and the 10x cell barcode (Fig. 1B). We performed PCR amplification and sequencing separately on the three batches of embryos. We then developed a custom analysis pipeline to extract both enhancer and cell barcodes from the sequencing reads, subsequently integrating the identified cell-enhancer pairs with the scRNA-seq dataset (see Methods).

Out of the 6,588 cells in our dataset, we identified 1,098 cells carrying an enhancer from the scRNA-seq libraries, and 910 cells through targeted PCR amplification. Notably, 349 cells were common to both methods, and 561 cells were only identified by targeted PCR, highlighting the value of PCR sequencing in corroborating and enriching our dataset. Importantly, these cells displayed no bias towards any specific tissue. Moreover, the cells detected by targeted PCR amplification were predominantly found in the same cell types where the enhancer had already been identified by scRNA-seq alone (Supplementary Fig. S16-S18), further validating the complementary nature of these two methodologies.

In conclusion, simply adding a short barcode to each candidate enhancer increased by 33% the number of cells carrying an active enhancer detected by spatial-scERA, significantly improving our ability to reconstruct each enhancers’ spatial activity.

### Spatial-scERA successfully captures enhancer activity in single cells

With a comprehensive understanding of the cell types in our dataset and an optimized protocol to efficiently capture cells containing an active enhancer, we proceeded to generate a global overview of the activity of the enhancers present in our library. Overall, 23 out of the 25 candidate enhancers were successfully retrieved in our combined dataset. Two enhancers were missing from the dataset (CRM5 and CRM8), suggesting that they are probably not active at stage 6. The number of cells per enhancer varied significantly, ranging from a single cell for psc_E14 to 320 cells for shg_A, with an average of 60 cells per enhancer. We analyzed the distribution of each enhancer across the different clusters by calculating the percentage of cells in which the enhancer is present within each cluster (Supplementary Table 2). Eleven enhancers were clearly active in the embryo, either across multiple clusters (for example GMR77A12, present in 294 cells and salm_blastoderm_early_enhancer, present in 58 cells), or in a tissue-specific manner (for example twi_ChIP-42, present in 128 cells, most of which are found in the mesoderm cluster (Fig. 2D, Supplementary Fig. S1-S13)). The remaining 12 enhancers were active in very few cells. While the GMR83E01 (15 cells) and CRM3 (19 cells) enhancers were enriched in the yolk cluster, no discernible patterns were immediately apparent for the other 11 enhancers (Fig. 2D, Supplementary Fig. S1-S13, Supplementary Table 2).

To verify whether the enhancers missing or weakly detected in the dataset were indeed not active at stage 6, we performed RT-qPCR in a subset of individually-generated reporter lines (Supplementary Fig. S19). We confirmed a complete absence of activity for CRM5 and CRM8, corroborating their absence in our spatial-scERA dataset. Similarly, CRM6 and CRM12, which were recovered in very few cells, exhibited a very low activity. Conversely, enhancers identified in many cells by spatial-scERA were also highly active in RT-qPCR. Overall, the results observed by RT-qPCR were consistent with the spatial-scERA data.

By integrating scRNA-seq and targeted PCR amplification, we identified potential tissue-specific activity for 11 out of 25 enhancers in our dataset. However, scRNA-seq only samples a small proportion of all the cells where the reporter is active in an embryo. While visualizing these few cells in a UMAP may sometimes be sufficient to predict the tissues where the enhancer is active, it is inadequate to determine the precise pattern of enhancer activity. To address this limitation, we used novoSpaRc, a computational tool capable of precisely mapping scRNA-seq data onto a virtual representation of any tissue or organism via optimal transport, in our case a *Drosophila* embryo.

### *In silico* reconstruction of stage 6 *Drosophila* embryos from spatial-scERA data with novoSpaRc

To reconstruct the spatial activity of the genes expressed in our scRNA-seq dataset *in silico*, we projected our sequenced cells onto a virtual stage 6 embryo composed of 3,039 positions using the novoSpaRc computational framework (26,27). NovoSpaRc uses optimal transport to infer a mapping between the gene expression space from a scRNA-seq dataset to the tissue space of the sample of interest, which is then used to predict the spatial expression of any given gene. NovoSpaRc spatial reconstruction accuracy is improved when used in conjunction with an atlas providing prior information on the spatial expression of informative marker genes. In this case, we took advantage of a pre-existing atlas consisting of 84 marker genes expressed at stage 6 (28), which recapitulates most expression patterns distinguishable during early *Drosophila* embryogenesis. We used this atlas to uncover the spatial position of the cells in our scRNA-seq dataset and plot their expression level. However, as the atlas does not account for internal cells, we first excluded yolk, hemocyte, and unknown clusters from our scRNA-seq dataset. The remaining 5,475 cells were re-analyzed from the raw matrix and resulted in 9 cell-types, including a novel cell-type labeled as head mesoderm (Fig. 3A-B).

**Fig. 3.**
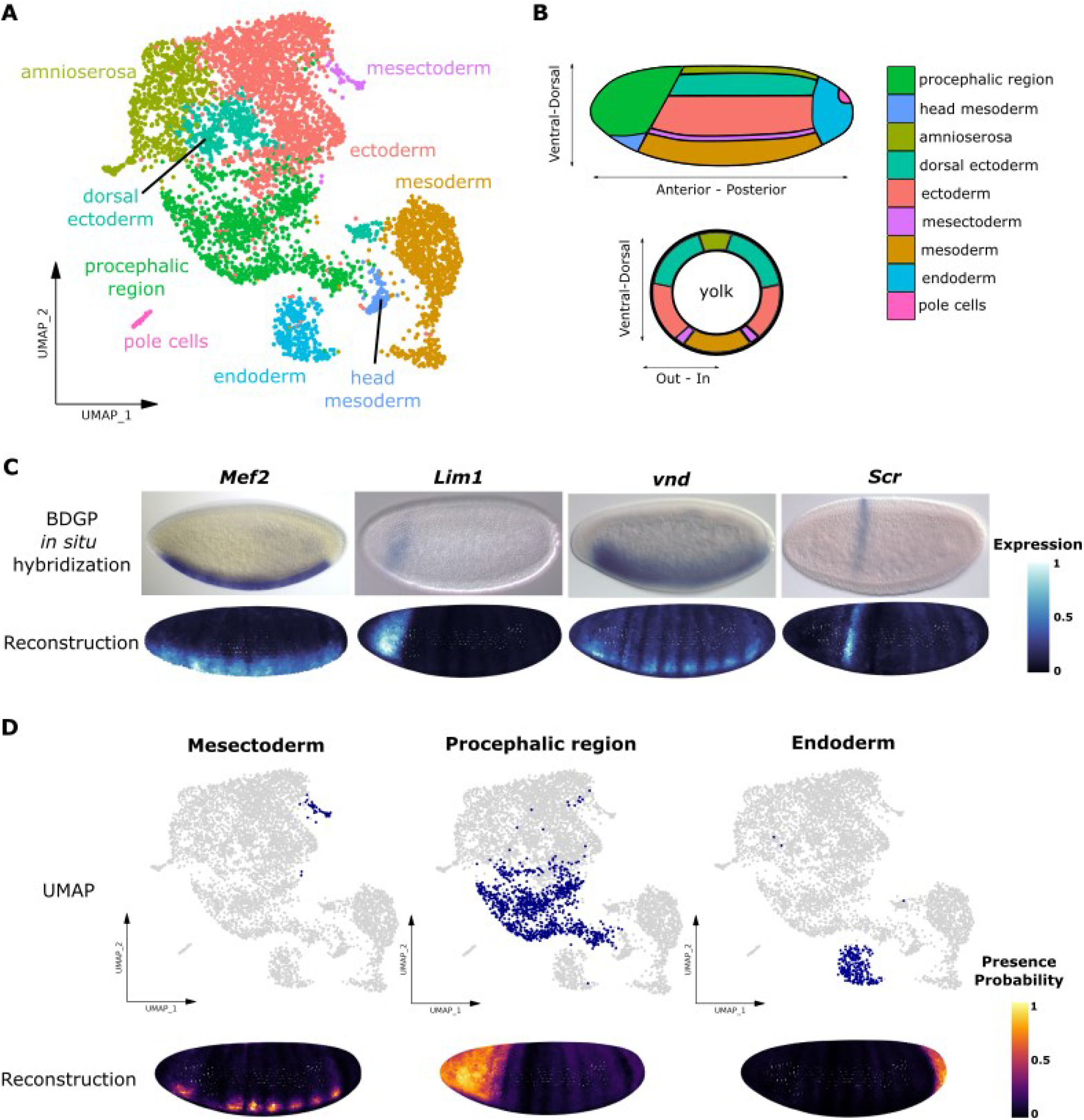
Validation of the spatial reconstruction. (**A**) UMAP of the reduced version (n=5,475 cells) of the scRNA-seq dataset with the 9 identified cell-types. (**B**) Schematic of a stage 6 *Drosophila* embryo displaying the expected location of the cell-types present in the scRNA-seq dataset. (**C**) Comparison between published *in situ* hybridization from the BDGP *in situ* database (top) (37) and spatial reconstruction with novoSpaRc (bottom) for 4 different genes. In the reconstruction, the lighter the blue, the higher is the level of expression. (**D**) Projection of the cells forming a cluster onto the reconstruction. UMAPs indicate in blue the cells forming the selected cluster. In the reconstruction, the color code indicates the probability of presence of the cells in the reconstruction from dark - low probability, to yellow - high probability.

We first validated the accuracy of the reconstructions by comparing them to the expression of well-known tissue-specific marker genes (Fig. 3C). As expected, cells expressing the mesodermal *Myocyte enhancer factor 2* (*Mef2*) gene were positioned ventrally. Similarly, we were able to accurately reconstruct the expression of the *LIM homeobox 1* (*Lim1*) and *ventral nervous system defective* (*vnd*) genes. Even genes with highly restricted expression, such as *Sex combs reduced* (*Scr*) which is expressed in a single stripe, were well reconstructed. However, we observed that the reconstructed expression pattern of some genes was sometimes slightly distorted, displaying stripe-like patterns. This is visible, for example, in the reconstructions of the *Mef2* and *vnd* genes (Fig. 3C), and is also seen in the reconstruction based on the *Drosophila* dataset associated with novoSpaRc (28) (Supplementary Fig. S20). This phenomenon might be due to a bias in the atlas with the over-representation of genes active in stripes.

To evaluate the statistical robustness of our approach, we assessed the accuracy of the reconstruction by comparing it to the expected pattern in the atlas using a leave-one-out cross-validation strategy. We used the Structural Similarity Index Measure (SSIM) as our evaluation metric (43,44), a tool originally developed for image analysis that reliably captures similarity based on overall intensity, the degree of intensity variation, and the preservation of spatial expression patterns. Our analysis yielded a mean SSIM score of 0.45 across the 84 genes in the atlas, with a distribution that differed significantly from a random model (p-value = 2e-107, KS-test; see Methods and Supplementary Fig. S21). These results indicate that the reconstruction generated from this atlas is statistically robust, enabling us to assess gene expression and enhancer activity with confidence.

Having established that novoSpaRc faithfully reconstructs the spatial expression of the genes in our dataset, we next applied this tool to reconstruct the expression of the *CD2* reporter gene for each candidate enhancer. Because spatial-scERA combines information from scRNA-seq with targeted PCR amplification, working with the expression level of the reporter is not possible anymore. Enhancer activity is thus not assessed based on reporter expression but by the presence or absence of an enhancer, identified either by its sequence (scRNA-seq) or by its barcode (targeted PCR) in each cell. To better visualize this binarized data, we decided to plot the probability of presence of the cells containing an active enhancer within the virtual embryo. To achieve this, we upgraded the novoSpaRc pipeline by allowing the software to plot the predicted location of a group of cells in the reconstruction. Positions where the highest number of cells are mapped will be assigned a higher probability of presence. As a result, the spatial reconstruction will display the probability of presence of the enhancer at any position of the embryo, even if the actual number of sequenced cells does not cover the full pattern. We tested this feature by mapping the spatial location of the same four genes used to validate our spatial reconstruction, confirming that the reconstruction based on the probability of presence correctly recapitulates the genes’ expected expression (Supplementary Fig. S22). We then applied this approach to map the spatial location of each of the nine scRNA-seq clusters onto the virtual embryo. Eight out of nine clusters were mapped to their expected spatial location (Fig. 3D, Supplementary Fig. S23), further validating this approach and the overall accuracy of the reconstruction.

### Spatial information must be considered for proper scRNA-seq cluster annotation

The ninth cluster, initially labeled as dorsal ectoderm, deviated from expectations. At stage 6, the dorsal ectoderm consists of a row of cells on the dorsal side of the embryo, between the amnioserosa and the lateral ectoderm (Fig. 4A). However, our spatial reconstruction mapped these cells along two distinct stripes at the opposite ends of the trunk region. To understand this discrepancy, we examined the actual expression pattern of the top differentially expressed genes in the dorsal ectoderm cluster. While “dorsal ectoderm” was the most common annotation for these genes in the BDGP database, many were in fact segmentation genes expressed in *anterior*-*posterior* stripes, such as *spalt major* (*salm*), *ken and barbie* (*ken*), and *Scr* (Fig. 4B, Supplementary Fig. S24). In contrast, *bona fide* dorsal ectoderm genes such as *zerknüllt* (*zen*) and *Dorsocross2* (*Doc2*) were not enriched in this cluster, confirming the inaccurate annotation (Fig. 4C).

**Fig. 4.**
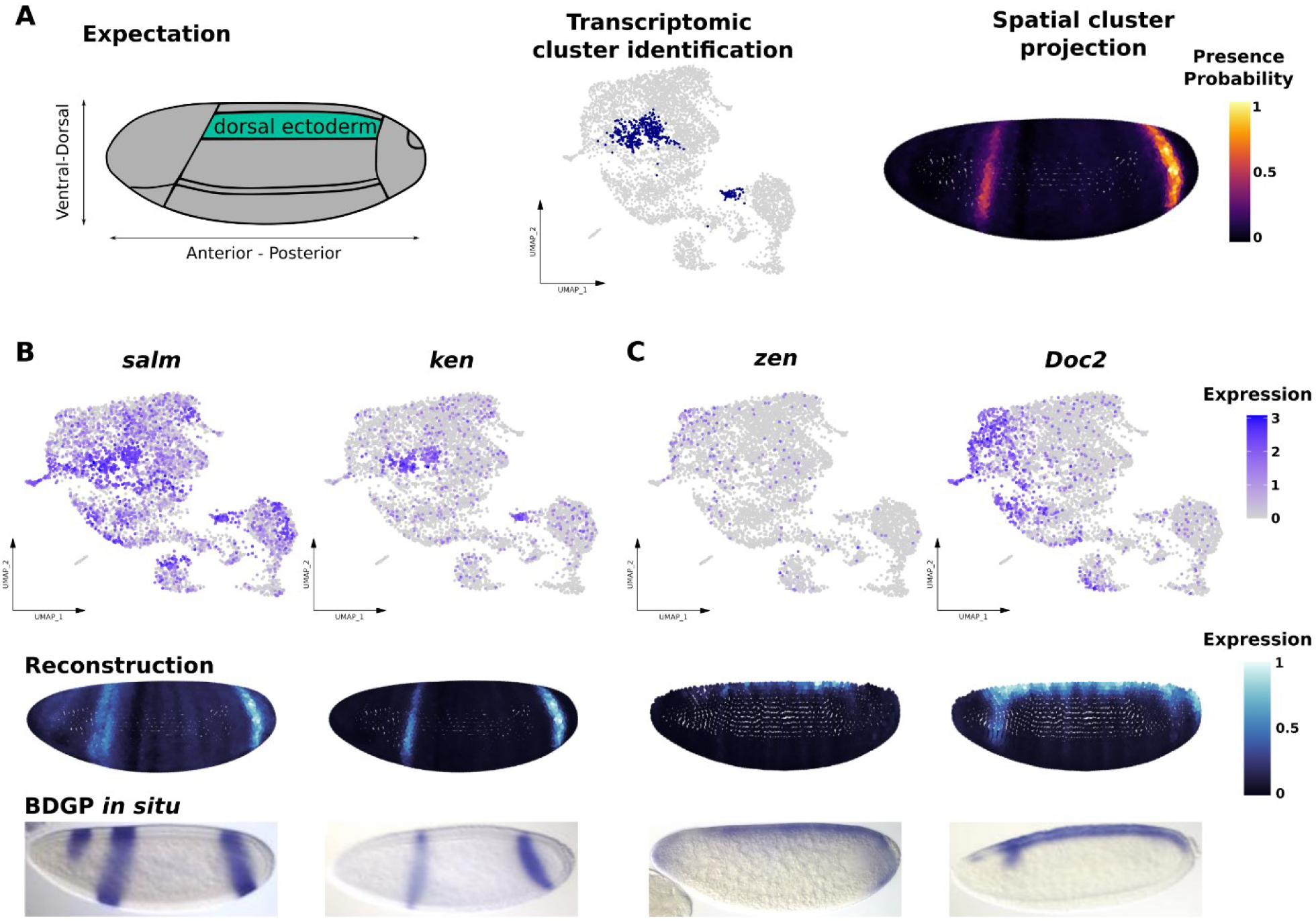
Spatialization identifies a mislabeled cluster. (**A**) Representation of the encountered mislabeling. Left: the location of the dorsal ectoderm in a stage 6 *Drosophila* embryo is highlighted in cyan. Middle: The location of the cluster annotated as dorsal ectoderm in the UMAP is highlighted in blue. Right: Location of these cells after projecting them on the virtual embryo. (**B-C**) UMAPs (top), reconstruction (middle) and published *in situ* hybridization from the BDGP *in situ* database (bottom) (37) of 2 genes differentially expressed in the mislabeled cluster (**B**) and 2 genes actually expressed in the dorsal ectoderm (**C**).

Having established that this cluster was not dorsal ectoderm, we then verified that our spatialization was able to correctly reconstruct the expression of the genes wrongly assigned to the dorsal ectoderm and of actual dorsal ectoderm genes. In both cases, the reconstruction accurately reflected the expression patterns observed by *in situ* hybridization, confirming that the mislabeling was due to the annotation of the cluster in the scRNA-seq data, rather than an issue with the reconstruction (Fig. 4B, C).

In conclusion, this result highlighted a key limitation of scRNA-seq: clusters do not necessarily correspond to distinct cell types but rather to sub-populations of cells with a similar transcriptome. Therefore, clusters cannot be accurately annotated based solely on *a priori* knowledge of a few highly differentially expressed genes. To ensure correct cluster labeling, transcriptomic information must be combined with spatial information.

### Spatial-scERA accurately confirms the spatial enhancer activity of known enhancers

Having established that our spatialization method faithfully reconstructs the spatial position of a group of cells in our dataset, we proceeded to predict the activity of the 25 candidate enhancers under study. For this purpose, we plotted the probability of presence of every cell where a given enhancer was identified by scRNA-seq or targeted PCR at each position of the virtual embryo. To validate our predictions, we generated stable fly lines for 22 out of the 25 enhancers, and analyzed the expression of the *CD2* reporter gene by *in situ* hybridization. We first focused on the five positive controls, comparing reconstruction results to previously published *in situ* hybridization experiments (33,35,45–48) or to our own images. We evaluated the reconstructions through visual inspection and numerical comparison of activity levels along the antero-posterior axis of the embryo, comparing the predicted activity patterns to the expected ones (Fig. 5A-D, Supplementary Fig. S1, S2, S3A). The results showed a strong correlation between the predicted reconstruction and the known pattern for all five control enhancers. For example, twi_ChIP-42 was correctly predicted to be active throughout the mesoderm and head mesoderm (33) (Fig. 5A). vnd_743 was accurately mapped to the mesectoderm and procephalic region (46) (Fig. 5B). The spatial reconstruction of salm_blastoderm_early_enhancer accurately predicted its activity in three main stripes in the procephalic, cephalic furrow and posterior trunk regions (45) (Fig. 5C). The spatial reconstruction also predicted activity in additional stripes in the trunk, which are faintly visible in our *in situ* hybridization experiments. The activity of the eve_late_variant enhancer was predicted to follow a stripe pattern. However, some of the middle stripes were missing, probably due to the very limited number of cells carrying this enhancer in our dataset (Supplementary Fig. S1A). The most striking result was observed with h_stripe1. Spatial-scERA identified this enhancer in just 9 cells scattered across the UMAP, making tissue-specific activity determination challenging from this data alone. However, the reconstruction revealed a precise stripe that perfectly matched the expected location (35) (Fig. 5D). Finally, as expected, the ChIP-27 enhancer which was used as a negative control did not display any activity in the reconstruction (Supplementary Fig. S3B). Overall, this confirmed that spatial-scERA can accurately predict enhancer activity, even with as few as 9 cells in the dataset.

**Fig. 5.**
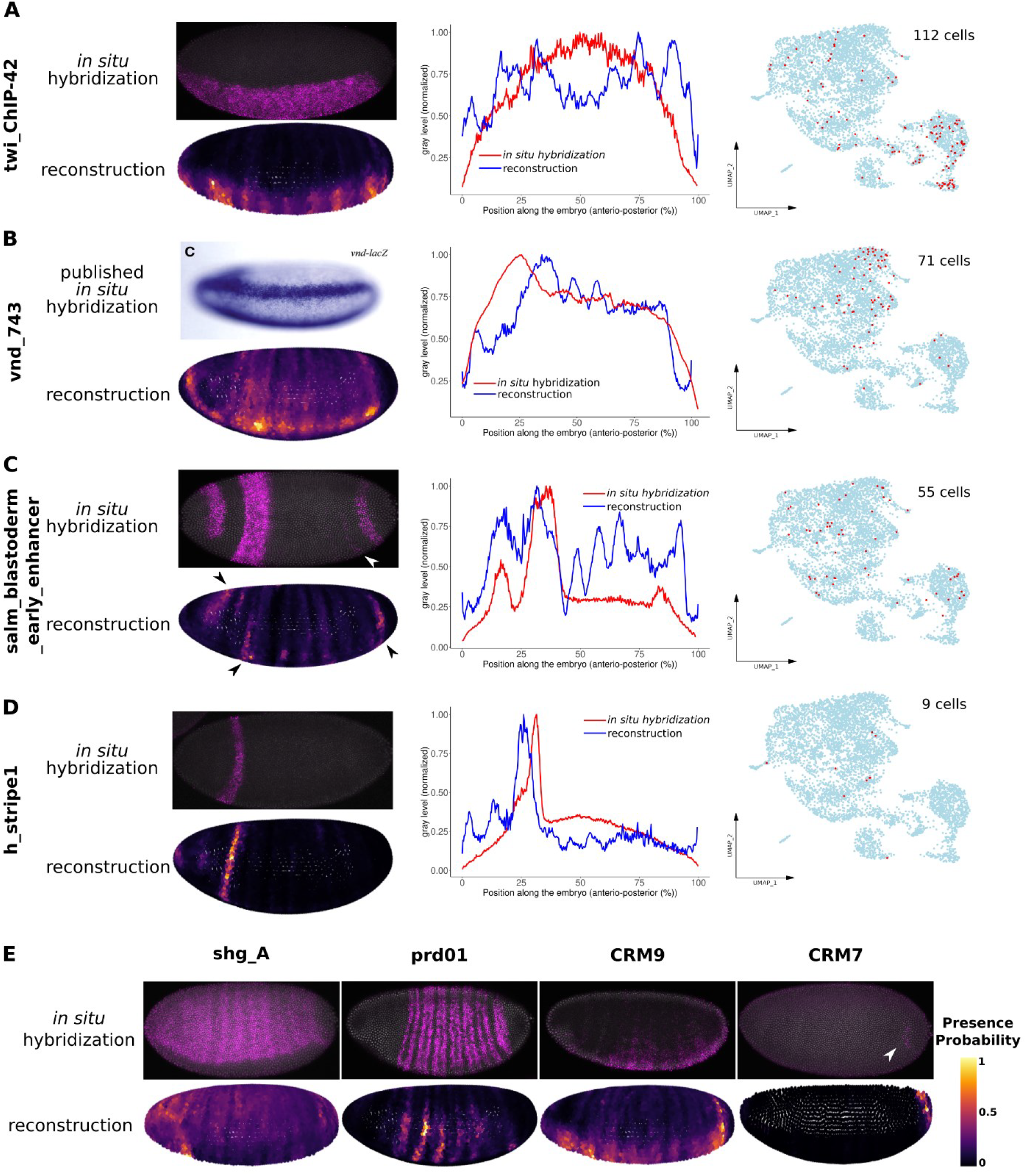
Spatial-scERA accurately predicts enhancer activity. (**A-D**) Left: Comparison of the activity observed by *in situ* hybridization and our spatial reconstructions for 4 control enhancers. Middle: Profiles of average expression levels across the entire embryo by *in situ* hybridization experiments (red) and in the reconstructions (blue). Right: Number of cells with active enhancer and their location in the UMAP (in red). Note: *in situ* hybridization data from the vnd_743 enhancer was obtained from (46). The embryo was imaged from a more ventral view than the reconstruction. (**E**) Examples of predicted spatial activity for 4 enhancers (bottom) in comparison to *in situ* hybridization (top). Faint expressions are pinpointed with white arrows.

### Spatial-scERA can be used to predict the spatial activity of uncharacterized regions

Having validated the accuracy of our enhancer predictions, we proceeded to generate reconstructions for the 19 uncharacterized putative enhancers in our dataset. As mentioned previously, CRM5 and CRM8 were not active in the scRNA-seq dataset, indicating that these enhancers are likely not active at stage 6, a conclusion confirmed by imaging the corresponding fly lines (Supplementary Fig. S9A, S10B). The psc_E14 enhancer was detected in only a single cell in the full scRNA-seq dataset, which was discarded in the reduced dataset used for reconstruction (Supplementary Fig. S4B). We therefore concluded that this enhancer was not active at stage 6. Among the other enhancers, shg_A, GMR77A12, GMR83E01, and CRM3 were identified in several cells of the yolk cluster. As the yolk was removed from the dataset for spatial reconstruction, we could not confirm this activity in the virtual embryo. However, activity in the yolk was confirmed by *in situ* hybridization, providing additional evidence that our spatial-scERA effectively captures cell-type specific enhancer activity (Supplementary Fig. S25).

Except for psc_E14, the other five enhancers known to be active at later stages were clearly also active at stage 6. prd01, shg_A, and GMR77A12 all displayed a relatively broad pattern: along 7 stripes within the lateral ectoderm for prd01 (Fig. 5E, Supplementary Fig. S4A), across nearly the entire embryo excluding the amnioserosa and trunk mesoderm for shg_A (Fig. 5E, Supplementary Fig. S5A), and throughout the trunk region for GMR77A12 (Supplementary Fig. S6A). GMR83E01 on the other hand displayed a more localized pattern in the procephalic region (Supplementary Fig. S6B). All these patterns were confirmed by imaging of the reporter lines (Fig. 5E, Supplementary Fig. S4-S6). However, the SoxN_5830 enhancer, predicted to be active only in the procephalic region, showed ubiquitous activity by imaging (Supplementary Fig. S5B). This discrepancy likely stems from the insufficient sampling of cells carrying the SoxN_5830 enhancer in the spatial-scERA dataset. Indeed, this enhancer was active in only 18 cells, which probably makes it difficult to predict such a broad pattern.

From the thirteen entirely uncharacterized candidate enhancers, some, such as CRM1, CRM3, CRM7 CRM9, CRM11, and CRM13 displayed clear but more or less extensive tissue-specific patterns, which were also confirmed by imaging (Fig. 5E, Supplementary Fig. S7A, S8A, S10A, S11A, S12A, S13A). However, some candidates were either not active (CRM5 and CRM8; Supplementary Fig. S9A, S10B) or predicted to be active in highly restricted patterns that were not confirmed by imaging (CRM6 and CRM12; Supplementary Fig. S9B, S12B). These often correspond to enhancers that were active in very few cells in the combined scRNA-seq and targeted PCR dataset. We therefore considered that when fewer than 10 cells were captured by spatial-scERA, the accuracy of the reconstruction should be treated with caution and ideally validated by other means. Conversely, CRM7, present in just 3 cells, was clearly predicted to be active along a ring of cells at the posterior end of the embryo, consistent with the faint signal observed by imaging (Fig. 5E, Supplementary Fig. S10A). Finally, the two discrepancies between the spatial-scERA predictions and our imaging data are in the activity of the CRM2 and CRM4 regions. For CRM2, the spatial reconstruction predicted a very localized activity at the posterior side of the dorsal region which is absent in our imaging data. Instead, imaging reveals a faint activity in the middle of the dorsal region (Supplementary Fig. S7B). No activity was detected by imaging CRM4, although spatial-scERA captured 26 cells with enhancer activity and predicted their spatial localization along antero-posterior stripes, partially recapitulating the expression of the closest gene *scabrous* (*sca*) (Supplementary Fig. S8B).

In conclusion, the comparison of reconstructed activity from spatial-scERA with imaging data confirmed that our method faithfully recapitulates the activity of candidate enhancers in 23 out of the 25 cases. These predictions span spatial activity patterns ranging from no or very weak activity to very broad patterns. This comparison highlights the value of spatial-scERA in identifying the spatial activity of enhancers within a multicellular organism.

### Spatial-scERA is a better predictor of enhancer activity than DNase I hypersensitivity or histone modifications

When comparing the enhancer activity reconstructions obtained with spatial-scERA to DNase I hypersensitivity and histone modification signal, we did not observe any obvious correlations. For example, regions such as psc_E14 which overlapped high DNAse I hypersensitivity and histone modification signal are in fact completely inactive in our spatial-scERA data (Supplementary Fig. S4B). Similarly, the ChIP-27 and CRM5 regions are inactive despite the presence of a DNAse I hypersensitivity peak (Supplementary Fig. S3B). Conversely, CRM10 and CRM13 are active despite the absence of a strong DNAse I hypersensitivity signal (Supplementary Fig. S11B, S13). Still, DNAse I hypersensitivity seems to be a slightly better predictor of enhancer activity than histone modifications. Indeed, CRM7 and CRM8 are located in closed chromatin regions and are not active in spatial-scERA despite strong peaks of H3K4m1 and H3K27ac histone modifications (Supplementary Fig. S10, S13). Conversely, salm_blastoderm_early_enhancer, vnd_743, and prd01 are broadly active enhancers despite the absence of a strong H3K27ac signal (Supplementary Fig. S2A, S3A, S4A). This demonstrates the value of spatial-scERA as a complementary tool to genome-wide enhancer prediction techniques (such as MPRAs or the mapping of DNase I hypersensitive sites and histone modifications) for the precise mapping of spatial enhancer activity in a multicellular sample.

### Spatial-scERA can be used to identify the enhancers’ putative target genes

Finally, we asked whether the spatial reconstructions generated by Spatial-scERA could be used to predict enhancer target genes and provide biological insights into enhancer-gene relationships. To this end, we used the SSIM score introduced earlier, systematically comparing the spatial reconstruction of each enhancer with that of all genes within a 600 kb window surrounding the enhancer. As a proof of principle, we first focused on the 11 enhancers in our dataset with known target genes. In nine out of eleven cases, the known target gene ranked among the top 10 genes with the highest SSIM scores (Supplementary Fig. 26A, Supplementary Table 3). Notably, for three enhancers (*vnd743, salm_blastoderm_early_enhancer*, and *GMR77A12*), the known target gene achieved the highest SSIM score. Conversely, two enhancers (*eve_late_variant* and *SoxN5380*) had particularly low SSIM scores, reflecting the poor quality of their spatial reconstructions due to insufficient cell sampling, as previously discussed. It is important to note that the SSIM score does not account for cases where an enhancer recapitulates only a subset of its target gene’s expression pattern. Nevertheless, in the case of the *h_stripe* enhancer, the *hairy* gene ranked fifth in SSIM score despite the fact that the enhancer is only active in a subset of the hairy expression pattern. To address this, we visually confirmed the overlap between enhancer activity and target gene expression by plotting both the enhancer reconstruction and the gene’s expression on the same virtual embryo (Supplementary Fig. 27).

Next, we applied this approach to ask whether the uncharacterized enhancers in our dataset regulate the expression of their nearest genes. While enhancers are often assumed to control the expression of their closest gene, in fact only 47% of enhancers interact with the nearest transcription start site (TSS) in the human genome (49). Even in the *Drosophila* genome, 73% of interactions span distances larger than 50 kb (50). To test this in our dataset, we used the SSIM score to compare the spatial activity of the CRMs to the expression of their nearest genes. Excluding CRM5 and CRM8 (which showed no detectable activity), we found that for CRM1, CRM4, CRM9, and CRM11, their nearest genes (*ImpL2, sca, sty*, and *Trim9*, respectively) ranked among the top 10 genes with the highest SSIM scores (Supplementary Fig. 26B, Supplementary Table 3). Visual confirmation of the overlap between CRM11 activity and *Trim9* expression further supported this result (Supplementary Fig. S27).

Interestingly, in some cases, the known or nearest gene was not necessarily the one with the highest SSIM score (Supplementary Table 3). For example, GMR83E01 is known to regulate *otp* at larval stages (51). However, the SSIM score of *otp* ranked eighth (SSIM = 0.10), whereas *Cht9* (SSIM = 0.48) and *CG15650* (SSIM = 0.27) ranked first and second, respectively (Supplementary Fig. 28A). Although the expression of these genes has not been characterized via *in situ* hybridization, their reconstructed expression patterns strongly resemble the enhancer’s activity (Supplementary Fig. S28A). Moreover, *Cht9* and *CG15650* are located approximately 161 kb and 70 kb from GMR83E01, across one or multiple TAD boundaries. Similarly, CRM2 does not recapitulate the expression of its closest gene, *vn*, but closely matches the expression of the *CG13288* (SSIM = 0.4424) and *CG32407* (SSIM = 0.6191) genes, located approximately 51 kb and 66 kb away across a TAD boundary (Supplementary Fig. 28B). An even more compelling example comes from the prd01 enhancer, which shares an overlapping expression pattern both with its target gene (*prd1*), with its closest genes, *firl* at 13 kb, and *CG14947* at 4 kb, and with *CG15480* (SSIM = 0.1108), located 1.1 Mb away across multiple TAD boundaries (Supplementary Fig. 28C). Using previously published Micro-C data from early *Drosophila* embryos (nuclear cycle 14) (52), we confirmed that the prd01 enhancer forms a long-range interaction with the *CG15480* locus, strongly suggesting a functional regulatory interaction.

Overall, these observations suggest that the GMR83E01, CRM2 and prd01 enhancers might be involved in a chromatin hub, regulating the expression of multiple genes, some of which are located across large distance and TAD boundaries. This is in line with previous findings demonstrating the presence of functional inter-TAD enhancer-promoter interactions in the *Drosophila* genome (34). Combining spatial-scERA with the SSIM score could thus offers a powerful framework to identify enhancer target genes and be highly complementary to chromatin conformation studies.

## DISCUSSION

We developed spatial-scERA to infer the spatiotemporal activity of candidate enhancer regions by projecting cells sequenced in a single-cell enhancer-reporter assay on a virtual reconstruction of the tissue of origin. Our method presents two key innovations. First, it combines scRNA-seq with targeted PCR to improve the identification of cells in which an enhancer is active. Second, it uses a custom version of novoSpaRc, a computational method for spatialization to reconstruct a map of enhancer activity on a virtual embryo. We applied spatial-scERA to 25 candidate enhancers in stage 6 *Drosophila* embryos and demonstrated that single-cell enhancer-reporter assays alone are not always sufficient to predict enhancer activity *in vivo*. In fact, we could establish the crucial requirement of spatialization for capturing complex spatially-defined expression patterns such as stripes. Overall, spatial-scERA faithfully recapitulated enhancer activity observed by imaging the same constructs in 23 out of 25 cases. Interestingly, 5 out of 25 enhancers were not active at stage 6 despite the presence of DNAse I and active chromatin modifications peaks indicating the opposite. Our work thus demonstrated that chromatin accessibility and enhancer-specific histone modifications alone are often poor predictors of enhancer activity (23). This highlights the importance of methods such as spatial-scERA, to capture precise spatiotemporal enhancer activity *in vivo* in multicellular organisms. Finally, we combined spatial-scERA with the SSIM score to predict enhancer target genes and identified potential regulatory long-range interactions.

A key feature of spatial-scERA is its ability to generate virtual enhancer activity maps. While many methods leverage the cellular resolution of scRNA-seq to enhance existing spatial transcriptomic data through deconvolution (44,49–52), there are far fewer tools available for reconstructing spatial data directly from scRNA-seq alone. Among these, different approaches are employed: Machine Learning (MLSpatial (53)), Optimal Transport (novoSpaRc (26)), or a combination of both (D-CE (54)). Although MLSpatial and D-CE are more recent developments, they have not demonstrated superior performance compared to novoSpaRc, especially when it is used in conjunction with an atlas. Additionally, novoSpaRc has been successfully applied to reconstruct a variety of distinct tissues, including the floral meristem (23 genes in the atlas (43)), human duodenum epithelium (22 genes in the atlas (55)), and human organoids (32 genes in the atlas (56)). Given that novoSpaRc remains the state-of-the-art tool for spatial reconstruction from scRNA-seq data, and considering that it had already been optimized for the *Drosophila* atlas, it was a logical choice for our study. While the reconstructions were overall of very high accuracy, we noticed a clear tendency to amplify stripe patterns in the reconstruction, probably due to the presence in the atlas of a majority of genes expressed in stripes (30 out of 84 genes). Moreover, when cells have a transcriptome that is too different from the one expected from the atlas, novoSpaRc tends to place them at locations where the atlas is less spatially informative. This highlights the need for more complex and extensive atlases. With the development of spatial transcriptomics technology such as multiplexed RNA imaging or spatially-resolved DNA sequencing (57), we believe that large atlases will become commonly available in multiple species. Our method could thus be extended to study enhancers in more complex tissues and any organism or organoid where an atlas is available.

The main limitation of single-cell enhancer-reporter assays remains the ability to capture and sequence the cells in which our enhancers are active. This can be particularly challenging for enhancers that are active in a very small portion of the sample. To circumvent this limitation, we have combined scRNAseq with targeted PCR amplification, hence greatly improving cell recovery. Moreover, thanks to our custom version of spatialization pipeline, we can faithfully reconstruct the activity of an enhancer by plotting its probability of presence at each position, even when as little as 9 positive cells have been sequenced. Nevertheless, accurate reconstruction still requires a minimum number of sequenced positive cells per enhancer. This number will vary depending on the number of cells in which the enhancer is active, but we could estimate, based on our comparisons with imaging data, that in most cases at least 10 cells are required to reconstruct the activity of a tissue-specific enhancer. Given this efficiency, we estimate that spatial-scERA can easily be scaled up to query at least 100 enhancers in a single experiment targeting 10,000 cells. While we present here a proof of concept with 25 candidate enhancer regions, we believe that with the decrease in cost and increase in efficiency of scRNA-seq experiments, it will become possible to routinely test a large set of candidate regions of interest and establish their spatial activity in an unbiased manner using spatial-scERA. This will ultimately pave the way for a more comprehensive characterization of *in vivo* enhancer biology in multicellular organisms.

## METHODS

### Candidate enhancer selection

The list of candidate enhancers was established by selecting non-coding DNA regions located in the vicinity (10kb up and downstream) of tissue-specific genes. The list of tissue-specific genes was generated using a scRNA-seq dataset from stage 6 *Drosophila* embryos (28). The expression matrix, consisting of 1,297 cells, was analyzed in R using the *Seurat* package (v4.4.0) (58). Highly variable genes were selected using the *mean*.*var*.*plot* method. Scaling and PCA were performed with default settings. The first 16 principal components were retained based on the knee observed with the *ElbowPlot* function. The *RunUMAP* function was used for dimensionality reduction and *DimPlot* for visualization. Clustering with *FindClusters* (resolution 0.5) resulted in six clusters. The 10 most differentially expressed genes for each cluster were identified using *findAllMarkers*, and formed the list of tissue-specific genes. Cluster identities were determined by retrieving the cell-type annotation of these genes from FlyBase (59) and the Berkeley *Drosophila* Genome Project *in situ* database (37).

These genomic loci were then visually inspected in a genome browser for the presence of the H3K4me1 and H3K27ac histone modifications based on ChIP-seq datasets generated in whole embryos at 0 to 4 hours after egg lay (29,30) and for the presence of open chromatin based on DNase I hypersensitivity in stage 5 whole embryos (15). We also verified that these regions were devoid of RNA-seq signal using a whole embryo dataset at 2 to 4 hours after egg lay (31). This resulted in 111 regions which we further narrowed down to 19 candidates by selecting the ones displaying higher level of DNase I hypersensitivity and/or histone modification and by cross-referencing them to *cis*-regulatory information available in the RedFly database (32). Six additional regions of interest to the team were added to this set, resulting in a total of 25 candidate enhancers. The exact coordinates of the regions were set to be centered around the DNase I hypersensitivity peak, with each region having an approximate length of 1000 bp (Supplementary Table 1). All tracks were plotted using pyGenomeTracks (v3.8) (60).

### Plasmid library preparation

The pBID-mphsp70-kozak-CD2-CmR/ccdB-25A-SV40polyA reporter plasmid (Supplementary Fig. 14) was generated from the pBID backbone vector (Addgene #35190, (61)) containing the *mini-white* gene as an integration marker. All plasmids were constructed using standard cloning methods with New England Biolabs restriction enzymes and T4 DNA ligase (New England Biolabs) or with the NEBuilder HiFi DNA Assembly kit (New England Biolabs). All constructs were verified by Sanger sequencing. The cassette containing the CmR and ccdB genes was amplified by PCR from the pSTARR-seq_fly-hsp70 plasmid (Addgene #71500, (18)). The other fragments of the final construct were synthesized by GeneArt (Life Technologies) and PCR amplified. All fragments were assembled into the pBID. The plasmid was propagated in ccdB survival bacteria (A10460, Life Technologies). The map of the final reporter vector is displayed in Supplementary Fig. S14. The reporter construct features the *hsp70* minimal promoter, a kozak sequence, the *CD2* gene followed by the pGL3’s SV40 late polyA signal. We also include a stretch of 25 adenosine upstream the polyA signal to avoid internal priming during the scRNA-seq library preparation (62) and ensure that most reads arise from the 3’ end of the transcript.

To prepare the library of 25 reporter plasmids, the candidate enhancers were amplified from genomic DNA of the w[1118]; PBac{y[+mDint2]=vas-Cas9}VK00027 (BDSC_51324) fly line. A 19 bp common sequence and a 6 bp specific barcode were added by PCR at the 3’ end of each candidate enhancers. The primers used for PCR are listed in Supplementary Table 4.

The 25 PCR reactions were gel purified and the candidate enhancer fragments diluted to 0.045 pmol/μl in a final volume of 8 μl. The fragments were mixed in equimolar ratio and inserted in batch in the pBID-mphsp70-kozak-CD2-CmR/ccdB-25A-SV40polyA plasmid, instead of the CmR/ccdB cassette, using AgeI and NotI restriction sites and the NEBuilder HiFi DNA Assembly kit (New England Biolabs). The vectors were transformed using Omni-max bacteria (C854003, Life Technologies), and directly transferred to 100 ml of LB medium containing ampicillin. The plasmid library was extracted using the NucleoBond Xtra Midi kit (740410.50, Macherey-Nagel).

### Injection and fly handling

The pool of 25 reporter vectors was injected in-house through PhiC31-mediated recombination (63). The injections were performed using a white-eyed fly line expressing the PhiC31 integrase and displaying a unique *attP* landing site on chromosome 2L (nos- ϕC31 / int.NLS; attP40 (64)). A total of 3,250 *Drosophila* embryos were injected with the reporter plasmid library. The resulting flies were crossed with the nos-ϕC31 / int.NLS; attP40 line. The 285 red-eyed transgenic progenitors were allowed to mate between themselves. The resulting flies are homozygous for the reporter construct but may contain a different candidate enhancer on each allele. These flies were amplified for three generations and used as a pool for single-cell experiments. We also extracted genomic DNA from a pool of transgenic progenitors to verify the correct genomic integration of the all 25 candidate enhancers by PCR and Sanger sequencing.

In parallel, we also derived homozygous lines carrying the same enhancer construct on both alleles for each candidate enhancer (with the exception of psc_E14, CRM10, and vnd_743) for further RT-qPCR and *in situ* hybridization experiments.

### Single-cell RNA sequencing

Freshly hatched adults from the pool of transgenic flies containing our library of candidate enhancers were combined in 5 embryo collection vials with standard apple cap plates. After three 45-minute pre-lays, *Drosophila* embryos were collected on apple juice plates for one-hour collection and incubated for another 2.5 hours at 25°C. We verified that the collected embryos mostly span developmental stages 5 to 7. The embryos were dechorionated using 2,6% bleach for 2 minutes, washed with water and PBS + 0.1% Triton X-100, finally resuspended in 1 ml of ice-cold PBS + 0.1% Triton X-100 and kept on ice.

Embryos were washed with ice-cold PBS, resuspended in 10 mL of dissociation buffer (PBS 1X + 0.1% BSA) and dissociated on ice in a Dounce homogenizer with gentle strokes of the loose pestle. This process was repeated for all collected embryos, using a small number of embryos each time. The cell suspension was transferred to 15 mL Falcon tubes and centrifuged for 10 minutes at 40 g at 4°C to pellet debris. Cells in the supernatant were transferred to new Falcon tubes and centrifuged again for 10 minutes at 800 g at 4°C. The supernatant was discarded, and cell pellets were combined and resuspended in 0.5 mL of dissociation buffer.

Two additional rounds of centrifugation, each for 5 minutes at 800 g at 4°C were performed to remove as much debris as possible.

Cells were counted using a Malassez counting chamber and the concentration was adjusted to 900 cells/μl in the dissociation buffer. scRNA-seq libraries were prepared using the Chromium Single Cell 3’ v3.1 protocol (10x Genomics), aiming for 10,000 cells per sample. Single cells were encapsulated into droplets in the Chromium Controller instrument for cell lysis and barcoded reverse transcription of mRNA. 40 μl of cDNA were recovered, and 10 μl (25%) were used for amplification, fragmentation, and Illumina library construction. The libraries were multiplexed and sequenced on a NovaSeq sequencer (Illumina) using 150-bp paired-end reads, yielding 500 million reads per library.

### Targeted PCR sequencing

From the remaining 30 μl of cDNA generated during the scRNA-seq library preparation, 300 ng were used for targeted PCR amplification. The region containing the specific enhancer barcode and the 10X Genomics barcodes (cellular barcode and UMI) was amplified with the Q5 Hot Start High Fidelity polymerase (New England Biolabs). The forward primer binds to the 19 nt sequence common to all constructs (GACGTCATCGTCCTGCAGG) and the reverse primer to the TruSeqRead1 sequence added during the scRNA-seq library preparation (CTACACGACGCTCTTCCGATC). The cycling conditions were as follows: Initial denaturation at 98°C for 10 seconds; then 25 cycles at 98°C for 10 seconds, 68°C for 30 seconds, 72°C for 20 seconds; then an elongation step at 72°C for 5 minutes.

The PCR product was purified using SPRIselect beads (Beckman Coulter) and 100 ng used to generate the final libraries using the NEBNext Ultra II DNA Library Prep Kit for Illumina (New England Biolabs). The libraries were indexed for multiplexing using NEBNext multiplex oligos kit for Illumina (New England Biolabs) and sequenced on a NovaSeq sequencer (Illumina) using 150-bp paired-end reads, yielding 10 million reads per sample.

### RT-qPCR

Freshly hatched adult flies of the appropriate genotype were placed in embryo collection cages with apple juice agar plates supplemented with yeast paste. After 3 pre-lay periods of 45 minutes for stage-specific collections, *Drosophila* embryos were collected for 1 hour and incubated for another 2.5 hours at 25°C until they reached stages 5-7.

The embryos used for RT-qPCR were directly transferred to the RA1 buffer supplemented with 2-Mercaptoethanol provided with the NucleoSpin RNA kit (Macherey-Nagel) and stored at -80°C for further RNA extraction.

RNA extraction was performed by grinding the embryos with a pestle in the RA1 buffer, followed by RNA purification using the Nucleospin RNA kit (Macherey-Nagel). Reverse transcription of 1 μg of RNA was performed using the RevertAid First Strand cDNA Synthesis Kit (Life technologies) with random primers. qPCR was performed with PowerUp™ SYBR™ Green Master Mix (Life technologies) for three independent biological replicates using the following primers:

- RpL32: ATGCTAAGCTGTCGCACAAATG and GTTCGATCCGTAACCGATGT
- CD2: CCTGAGAGCACCGTTTAAGT and AGATAGAGGGGCAGACCTTT

Relative quantification was done using the 2^−ΔΔCt^ formula.

### HCR RNA fluorescent *in situ* hybridization (FISH) in *Drosophila* embryos

Freshly hatched adult flies of the appropriate genotype were placed in embryo collection cages with apple juice agar plates supplemented with yeast. After 3 pre-lay periods of 45 min for stage-specific collections, *Drosophila* embryos were collected for 1 hour and incubated for another 2.5 hours at 25°C until they reached stages 5-7.

HCR RNA-FISH was performed according to the manufacturer instructions (Molecular Instruments) for whole-mount fruit fly embryos. All HCR probes, amplifiers and buffers were purchased from Molecular Instruments. Two different probe sets were designed; one against the *CD2* mRNA used with amplifier B3-488 to image spatial enhancer activity and one against *twist* mRNA used with amplifier B1-546 to help in embryo staging.

Embryos were mounted in ProLong Gold antifade reagent with DAPI (P36935, Life Technologies) and imaged on a Leica SP8 confocal microscope using a 20× glycerol objective. For each genotype, around 10 embryos were imaged and several Z-stacks were acquired (section thickness of 1.5 μm).

### Generation of a custom reference genom

Manual modifications were made to the *Drosophila melanogaster* genome from NCBI (GCF_000001215.4 RefSeq assembly) to integrate the reporter construct and enhancer sequences. We added extra chromosomes that contain the plasmid sequence and the sequence of each of the 25 enhancers (including the barcode). The plasmid was designated as chr5, and enhancer sequences were labeled chr6 to chr30, each with its own chromosome. A 100-bp flanking sequence was added to each enhancer on either side of the insertion site within the plasmid vector to improve the ability to map reads to the candidate enhancers.

The annotation file was manually edited to include an *mt* tag for genes on the mitochondrial chromosome. The *gene_id* and *gene_name* columns were swapped to prioritize *gene_name* during *Seurat* analysis. All features from the plasmid and enhancer chromosomes were categorized as *exon* in the gtf file to allow *CellRanger* to map reads at these loci.

The modified fasta genome and gtf annotation files were used as inputs for the *cellranger mkref* function to generate an index for mapping the fastq files.

### scRNA-seq analysis

Sequencing reads were controlled first by fastQC (65) to assure their good quality before mapping. They were aligned to the custom genome index using *cellranger count* (10x Genomics Cell Ranger v7.2.0) with 14 processing cores. The 10X Genomics cell and UMI barcodes were isolated by selecting the first 28 nucleotides in R1 with the parameter *r1-length* set to 28. After mapping, 24,886, 23,125, and 23,437 cells were obtained for replicates 1 to 3, respectively.

Each gene expression matrix went through stringent quality filtering. Non-viable cells, debris, or empty droplets were excluded if they contained fewer than 500 expressed genes and/or 2,000 UMIs. To prevent duplicates, a maximum threshold of 6,000 genes and/or 50,000 UMIs was set. Cells with more than 15% of mitochondrial genes or more than 36% of ribosomal genes were excluded. After quality control, 1,934, 1,963, and 2,691 cells were retained for each replicate.

To merge all three replicates, the *Seurat* V4 (58) integration vignette was followed. Datasets were normalized separately, and highly variable genes were identified using the *FindVariableFeatures* function with the *mvp* method. Both *FindIntegrationAnchors* and *IntegrateData* functions were used to generate a single dataset of 6,588 cells.

The standard Seurat pipeline was followed for analysis, including scaling, PCA, neighbor graph generation, clustering, and UMAP visualization. The first 30 components were selected for the PCA, and a clustering resolution of 0.5 yielded 16 clusters. Marker genes for each cluster were identified using *FindAllMarkers* with default parameters. To determine cell types, the top 10 genes with the greatest differential expression were analyzed using FlyBase (59) and the Berkeley Drosophila Genome Project *in situ* database (37). The most dominant annotation for the 10 genes was selected as the cluster annotation. If two clusters ended up with the same annotation, they were merged in one single cluster.

The reduced clustering was made by removing cells from the “unknown,” “yolk,” and “hemocytes” clusters using the *subset* function. The remaining 5,475 cells underwent the same analysis pipeline, resulting in 16 clusters, with 9 tissues visualized in two dimensions using UMAP.

*DimPlots, FeaturePlots*, and *Clustered DotPlots* were generated with the *scCustomize R* library (v2.1.2) to enhance Seurat visualization functions (66).

### Targeted PCR analysis

*Cutadapt* (v4.5) (67) was used to trim PCR-specific reads with a three-step script. Each step applied two trimming parameters: a minimal overlap of 10 nucleotides (*-O 10*) and removal of both reads if neither meet criteria (*--pair-filter=any*). Cutadapt allows a default error rate of 1 in 10 nucleotides.

Step one involved removing the 19bp common sequence (GACGTCATCGTCCTGCAGG) from forward reads and the 10X adaptor sequence (CTACACGACGCTCTTCCGATCT) from reverse reads. For the second step, both sequences were also checked on the opposite strand. All reads passing any of these two steps were merged into one forward (R1) and one reverse (R2) file. The final trimming removed the second common sequence (GCTGCCGCTTCGAGCAGACATGCATATG) and everything downstream, ensuring R1 reads contained only the 6-nucleotide enhancer barcode, while R2 reads retained only the first 28 nucleotides with the 10X cell+UMI barcodes.

In the specific case where the 10X adaptor sequence had been removed by Illumina’s trimming software, we used *seqkit* (v2.6.1) (68) to scan for 20 thymines within the 29th to 60th nucleotides, which is the region where the polyT sequence can be found if the adaptor was absent. For every reverse read in this case, if the forward read had the common sequence trimmed beforehand (GACGTCATCGTCCTGCAGG), the pair of reads was kept and sent back to the last trimming step.

We wrote a Python script to identify which enhancer was present within each cell. We observed that some of our reads contained enhancer barcodes that were different from the list of 25 barcodes used in the experiment, suggesting potential PCR amplification or sequencing errors. To correct for this, only reads containing one of the 25 expected enhancer barcodes and a cell barcode identical to one of the cells analyzed in the scRNA-seq analysis were kept.

For nearly every cell, more than 2 different enhancer barcodes were found, suggesting again PCR and sequencing error leading to the miss identification of our barcodes. To reduce the noise generated by such errors, a threshold strategy was used to detect the “true” enhancer in each cell. First, the frequency of all enhancer barcodes found in each cell was computed. If the most frequent enhancer barcode was found 10 times more often than the second most frequent, it was considered the real enhancer for that cell. If a cell had only one enhancer barcode, it needed to be present in at least 5 different reads to be considered real.

### Virtual embryo reconstruction using novoSpaRc

The expression matrix post *Seurat* analysis was loaded and processed using the *annData* python package (v 0.10.2) (69). Additionally, the atlas, representing one half of the surface of a stage 6 *Drosophila* embryo, was sourced from the novoSpaRc GitHub repository (28). This atlas comprises 3039 positions and captures the expression pattern of 84 marker genes.

The dataset was prepared by normalizing and converting the expression values into logarithmic values. Two cost matrices were computed to set up the reconstruction with parameters *num_neighbors_s* and *num_neighbors_t* set to 3 and 5, respectively. The reconstruction itself was executed with default parameters, except for the *alpha_linear* parameter, which was set to 0.35.

The output matrix, containing the probabilities of assigning each scRNA-seq cell to each position, was used to create two types of plots: a gene expression visualization on the reconstruction, and the distribution of the probability of presence of a defined group of cells on the reconstruction.

To evaluate the robustness of the reconstruction, a leave-one-out cross-validation was performed. For each gene in the atlas, we compared its original expression pattern with its reconstructed expression pattern after being omitted from the atlas. We manually encoded the SSIM equation to observe the similarity between the reconstructions and the atlas. For each gene, we randomized the reconstruction, keeping the same expression values but placing them randomly across the 3,039 positions, 100 times with a different seed. We used Kolmogorov–Smirnov test (KStest) to compare the similarity between the SSIM score of each gene expression pattern reconstructed after the leave-one-out strategy to the one of the random reconstructions.

To facilitate the generation of 3D interactive plots of the reconstruction, modifications were made to the core *pl*.*embedding* function to enable compatibility with *Plotly* (v 5.18.0) instead of *Matplotlib*. This involved adding a z-axis to the output matrix and implementing a *Matplotlib* normalization step within the loop that extracts values from the reconstruction. The reconstructions were visualized with *Plotly* by creating a 3D scatter of all locations and adding a second trace showing the expression level of a query gene on the reconstruction. Additionally, parameters such as a *threshold* parameter for displaying only positions with expression levels above a specified threshold and a *screenshot* parameter for automatically capturing dorsal, ventral, and lateral views were incorporated into the function.

Two additional functions were implemented to compute the probability of presence of multiple cells across the virtual embryo. The first function finds the index of a list of cells within the expression matrix, while the second function sums the presence probabilities of a list of cells over all positions before adding this “sum” column to the reconstruction object. Finally, a part of the script is dedicated to the creation of a table that recapitulates the list of cells that have each enhancer in the scRNA-seq and in the PCR, and merges them together into one final list per enhancer. These functions were used to plot the probability of presence for all the candidate enhancers, and for the cells coming from the same cluster.

### Comparing spatial-scERA reconstructions to HCR acquisitions

The RNA *in situ* hybridizations performed in this study by HCR, previously published *in situ* hybridization experiments, and spatial reconstructions were processed using *Fiji* (v1.54j) (71). Images were converted to 16-bit, background set to black, and intensity scaled from black to white.

To compare intensity levels, all embryos were standardized to the same length scale (0 for the anterior end and 100 for the posterior end). A rectangular region of interest was drawn around each embryo, and the plot profile function in *Fiji* was used to get the gray level across the embryo. Intensity values were then transferred to *R*, normalized between 0 and 1 based on the maximum intensity for each image, and curves were created using *ggplot2* (72).

### Identification of Putative Target Genes of Enhancers Using SSIM Scores

For each enhancer, we extracted all genes within a 600 kb genomic window using the *pybedtools* python library (v 0.10.0) (78,79). A custom Python script was developed to calculate the SSIM score comparing the enhancer activity and the expression of each gene in this region. Additionally, the genomic distance between the enhancer midpoint and the transcription start site (TSS) of each gene was computed. The SSIM scores for the top 10 genes with the highest SSIM score are listed in Supplementary Table 3. SSIM scores for the enhancer-target/closest genes were transferred to R for visualization as bar plots using *ggplot2* (77).

To identify potential new target genes, we leveraged a micro-C dataset from nuclear cycle 14 *Drosophila* embryos (52) at 5 kb and 1 kb resolutions. We created virtual 4C bedgraph files to visualize interaction scores between the enhancer regions and their surrounding genome using the *hicPlotViewpoint* function from the *HiCExplorer* package (v 3.7.2) (80). The virtual 4C data were plotted using *pyGenomeTracks* (v 3.8) (66).

To overlap the enhancer and target/closest gene spatial patterns, a custom novoSpaRc function was implemented. This function visualizes the gene expression positions on the reconstruction, overlays the enhancer activity, and highlights their co-expressed positions as a final layer. A threshold for expression values is adjustable through the “threshold” parameter for improved clarity in the visualization.

## DATA AVAILABILITY

All raw data were submitted to ArrayExpress (https://www.ebi.ac.uk/arrayexpress/browse.html) under accession numbers: E-MTAB-14447 (scRNA-seq), and E-MTAB-14445 (targeted PCR amplification). The custom dm6 *Drosophila* genome used for mapping with CellRanger can be downloaded on Zenodo (https://zenodo.org/records/14006160). The following publicly available databases and datasets were used: FlyBase r6.40 (https://flybase.org/) using the dm6 reference genome, scRNA-seq (GSE95025); DNase I hypersensitivity (SRA:SRX020691, SRA:SRX020692); histone modifications (GSE6273); RNA-seq (SRR1197368, SRR767626, SRR1197336); Micro-C (GEO:GSE171396); Redfly website (http://redfly.ccr.buffalo.edu/).

## CODE AVAILABILITY

Code and scripts used for analyses have been deposited on GitLab at https://gitbio.ens-lyon.fr/igfl/ghavi-helm/spatial_scera.

## ACKNOWLEDGMENTS

We are grateful to Laurent Gilquin and Sandrine Hughes for helpful advice in the design and analysis of targeted PCR sequencing data. We thank Sergio Sarnataro for all comments and contributions to the analysis of the scRNA-seq datasets. We are very grateful to Olivier Gandrillon and Laura Cantini for critically reading the manuscript. We thank all members of the Ghavi-Helm lab for discussions and comments on the manuscript. We also thank all the interns that made small contributions to the project throughout the years, in particular Nicolas Vaganay, Nathan Lecouvreur, and Louise Maillard. This work was technically supported by the IGFL sequencing facility (PSI), the IGFL microscopy facility, and the Arthro-tools facility of the Lyon SFR Biosciences (UAR3444/US8). This work was financially supported by an FRM starting grant (AJE20161236686) and ERC starting grant Enhancer3D (759708) to Y.G-H., by the EquipEx+ Spatial-Cell-ID under the “Investissements d’avenir” program (ANR-21-ESRE-00016), by a doctoral fellowship of the IADoc@UdL program to B.A., and by an FRM postdoctoral fellowship (SPF201909009228) to I.S.

## AUTHOR INFORMATION

### Affiliations

Institut de Génomique Fonctionnelle de Lyon, Univ Lyon, CNRS UMR 5242, Ecole Normale Supérieure de Lyon, Université Claude Bernard Lyon 1, 46 allée d’Italie F-69364 Lyon, France Aix-Marseille Université, MMG, Inserm U1251, Turing Centre for Living systems, Marseille, France

### Contributions

Y.G-H. conceived and supervised the study. P.V. co-supervised the computational aspects of the study. S.V., I.S. and Y.G-H. designed experiments, B.A., I.S., P.V., and Y.G-H. designed bioinformatics analysis. S.V. performed all experiments except microinjections which were performed by H.T. and D.L. B.A. assisted in the scRNA-seq experiment and performed all bioinformatics analysis. All of the authors discussed the results and implications and commented on the manuscript at all stages. Y.G-H. acquired funding.

