## Supplementary Figures for "Spatial-scERA: A method for reconstructing spatial single-cell enhancer activity in multicellular organisms"

**Supplementary Data**

Supplementary Figure 1

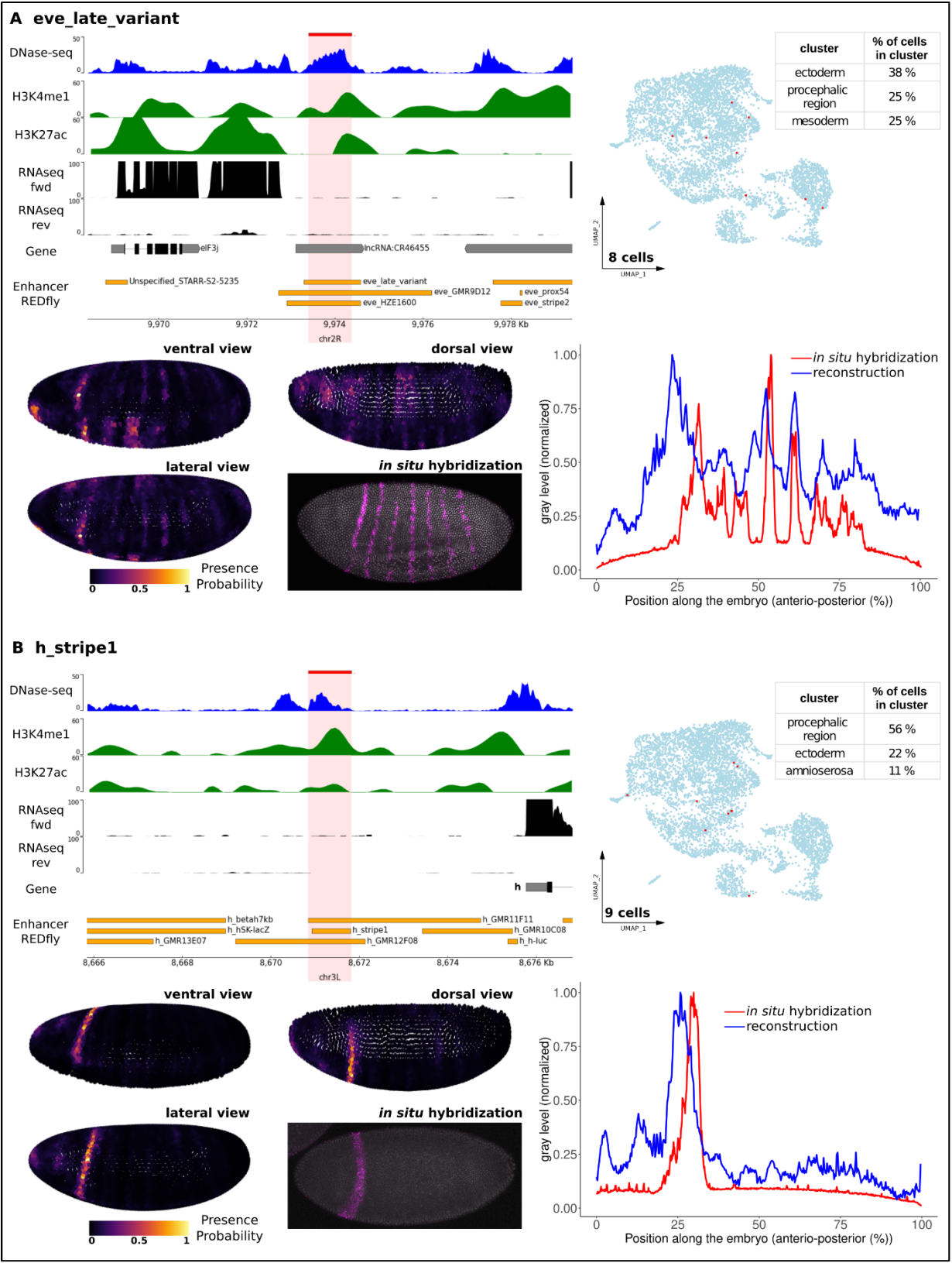

Supplementary Figure 2

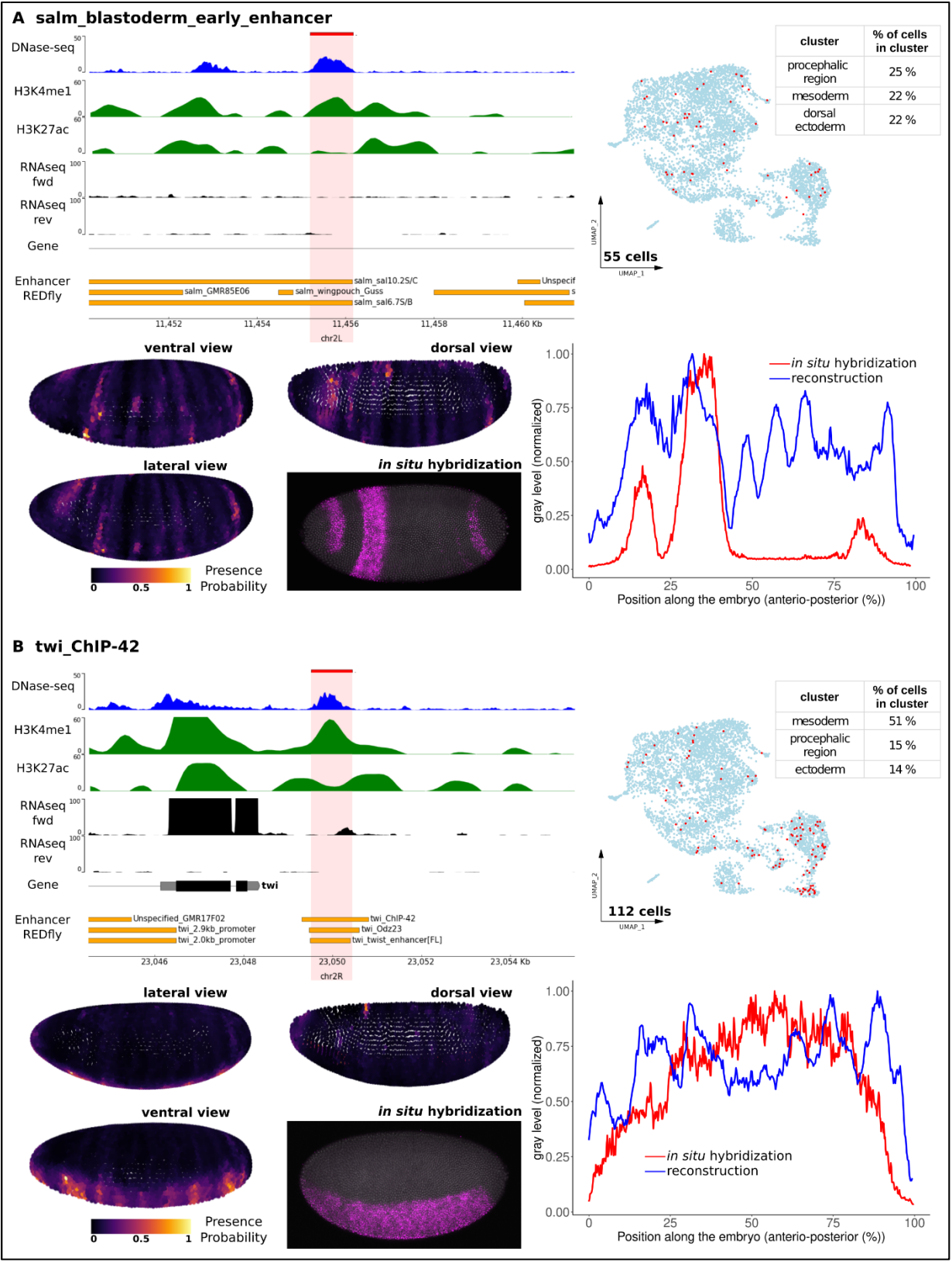

Supplementary Figure 3

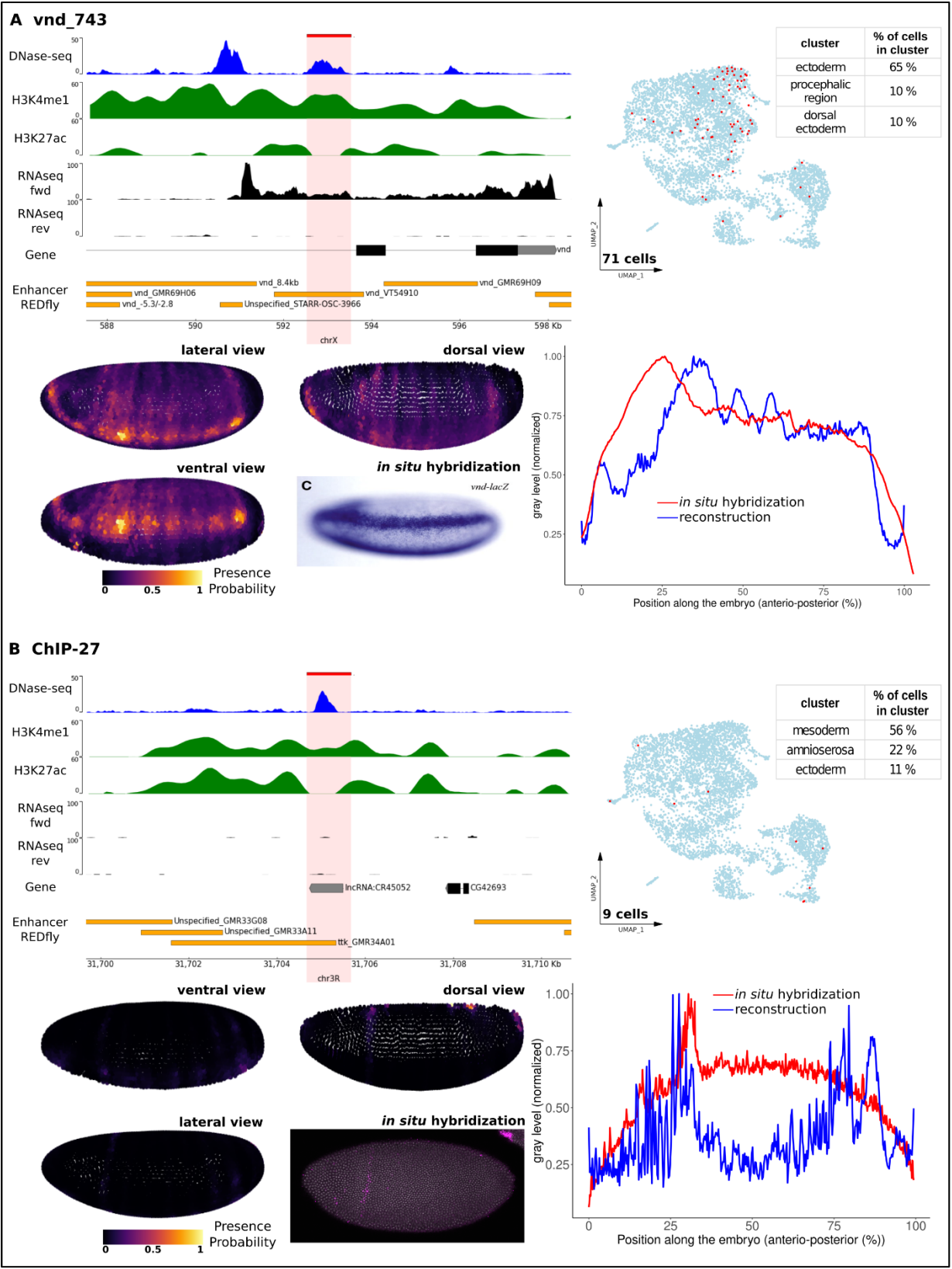

Supplementary Figure 4

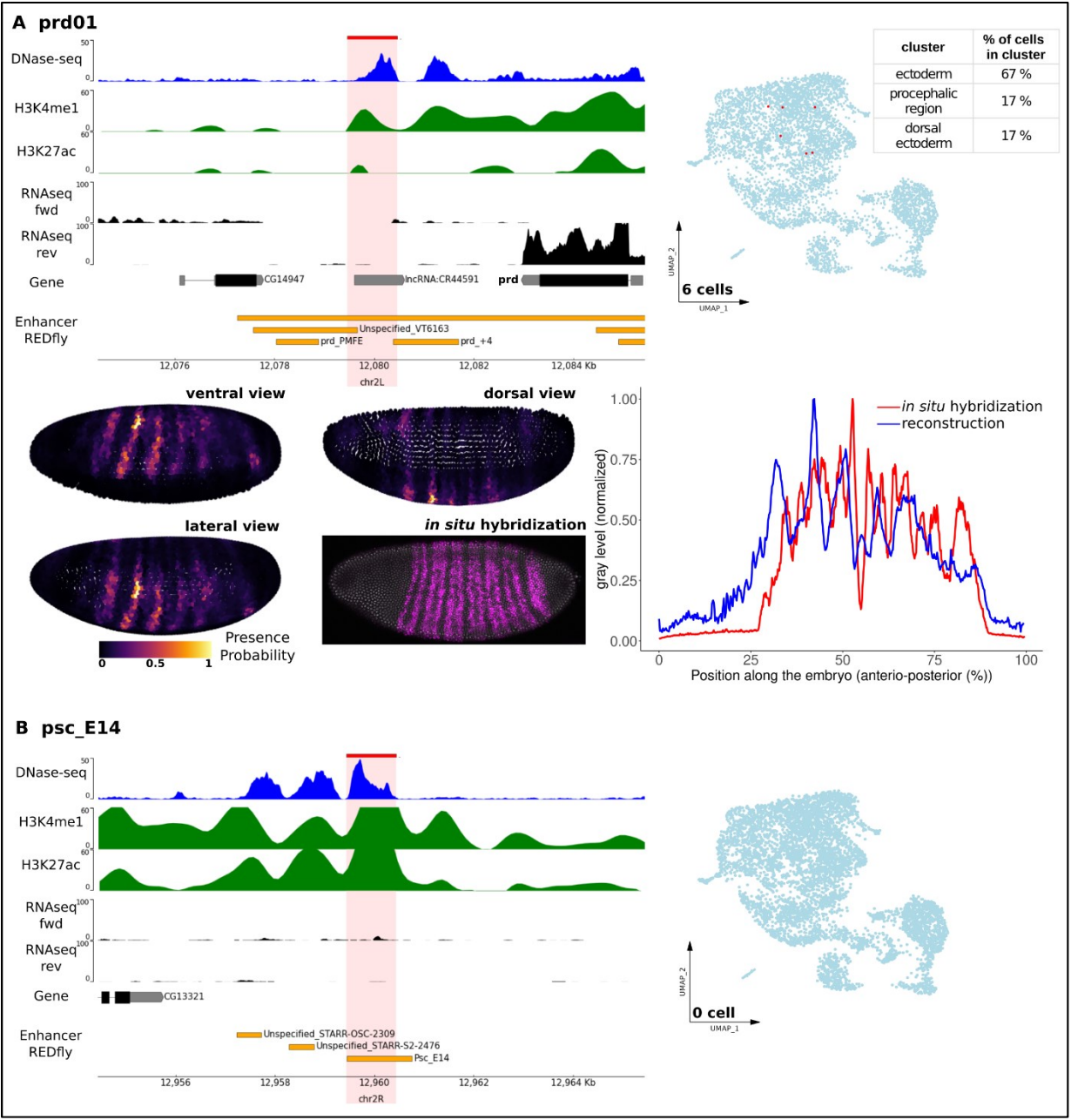

Supplementary Figure 5

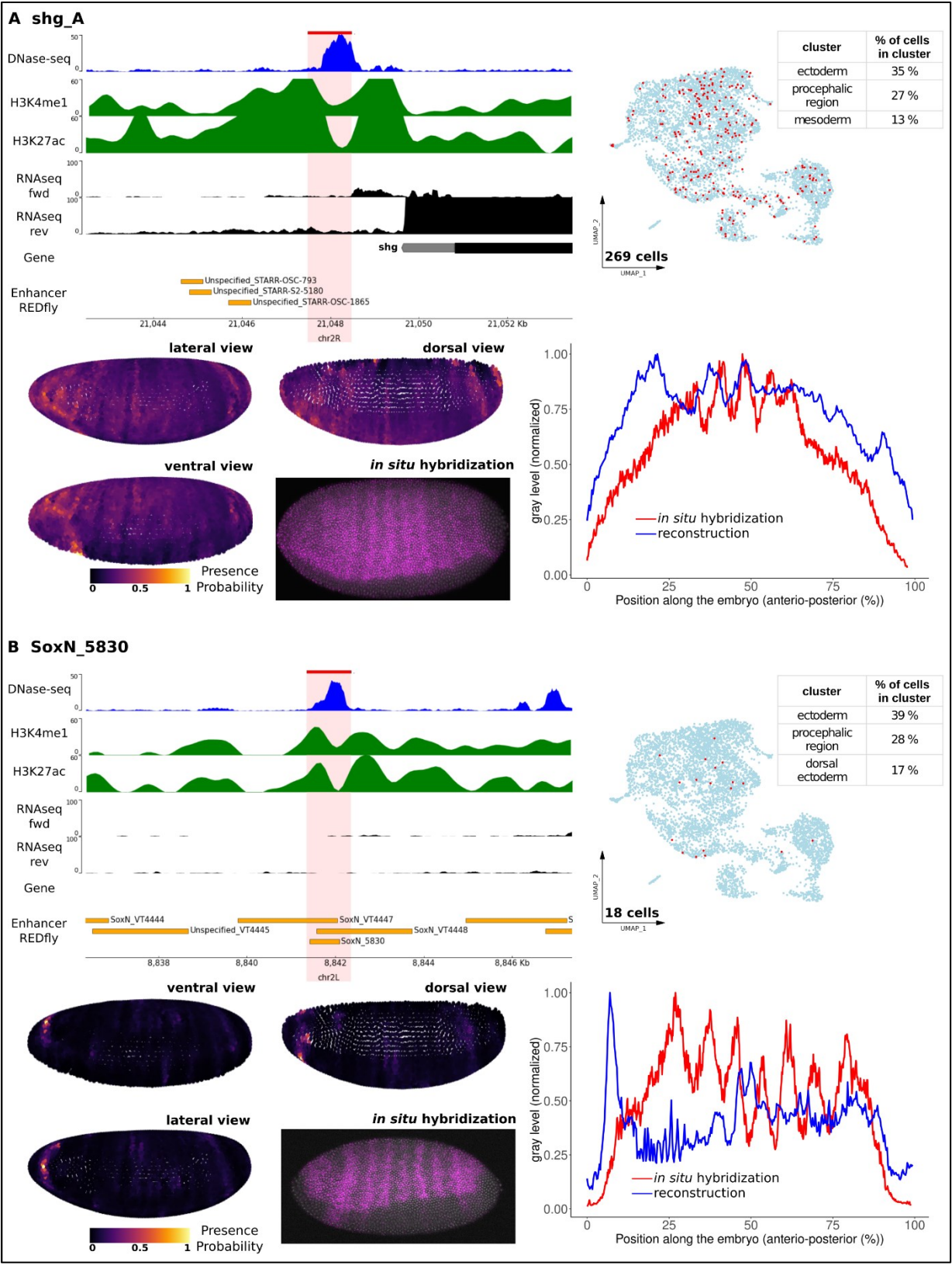

Supplementary Figure 6

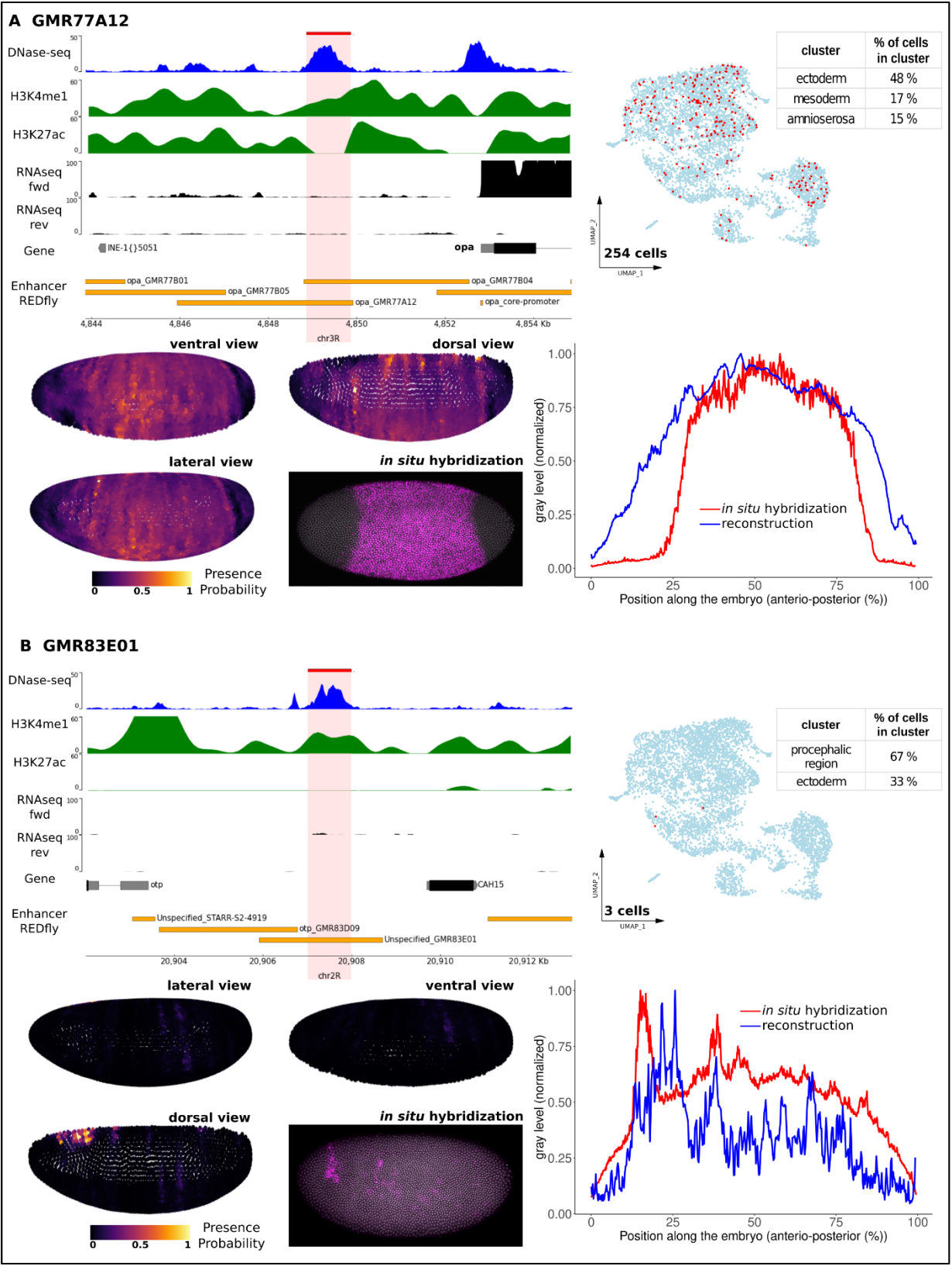

Supplementary Figure 7

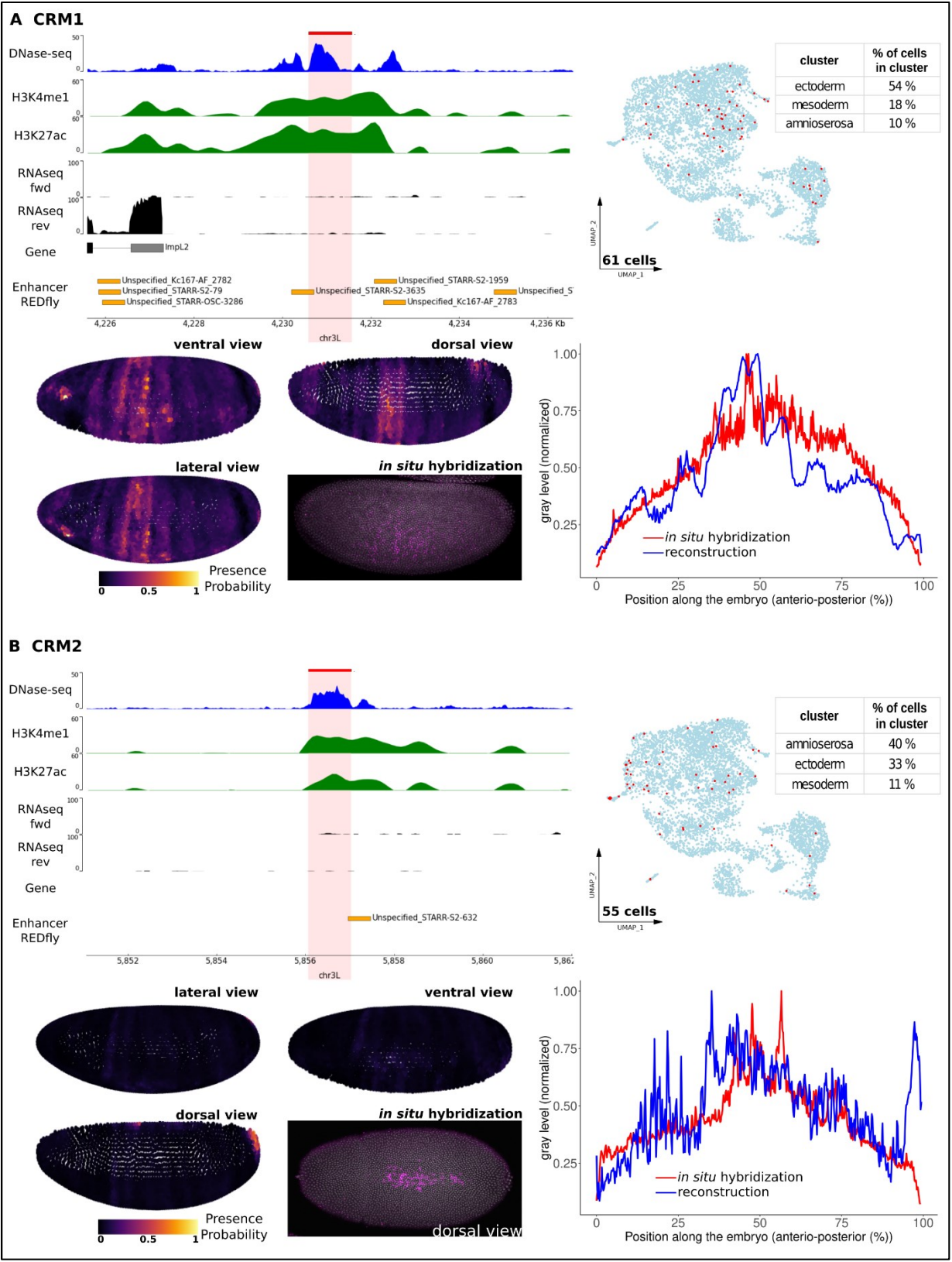

Supplementary Figure 8

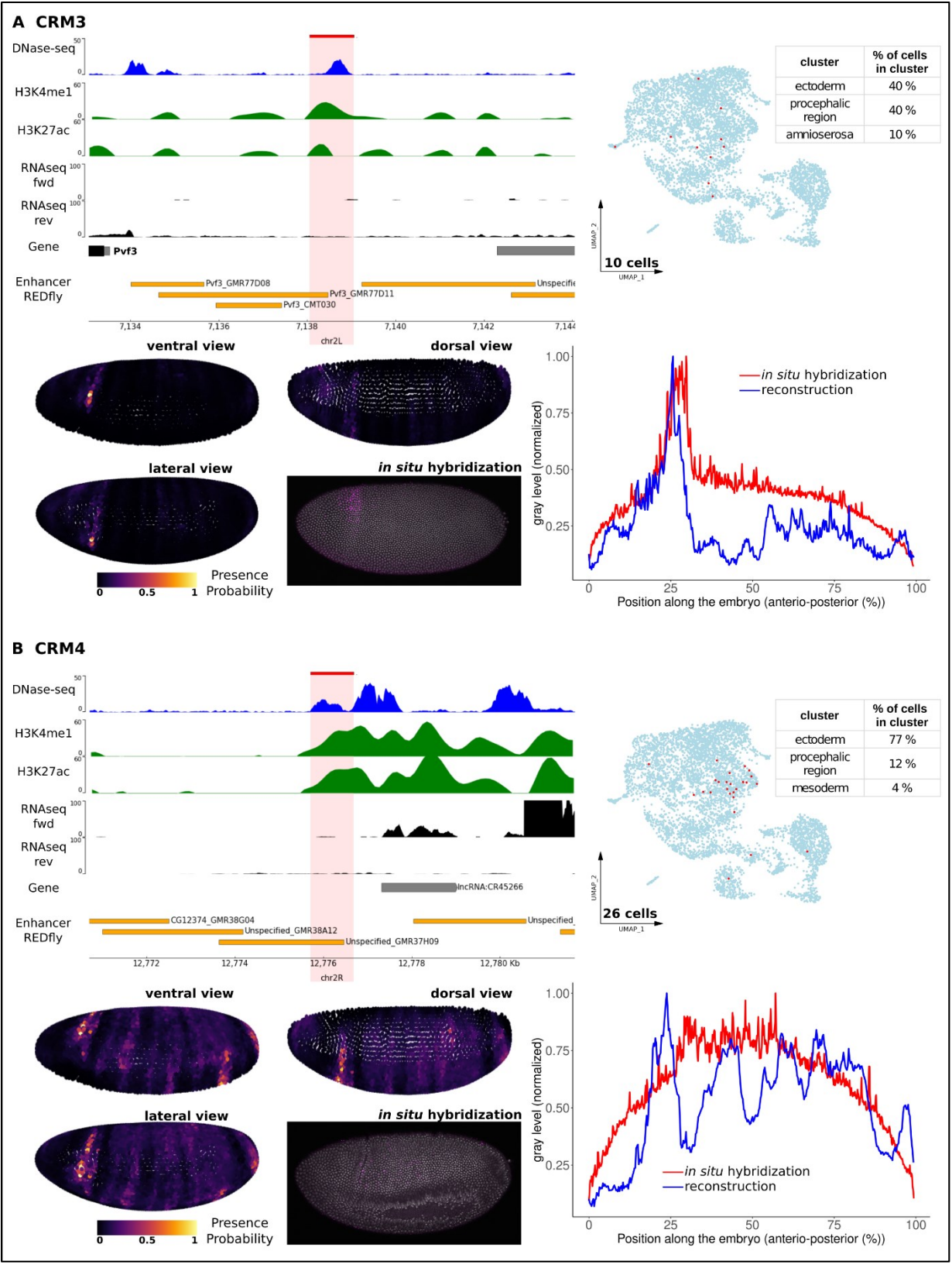

Supplementary Figure 9

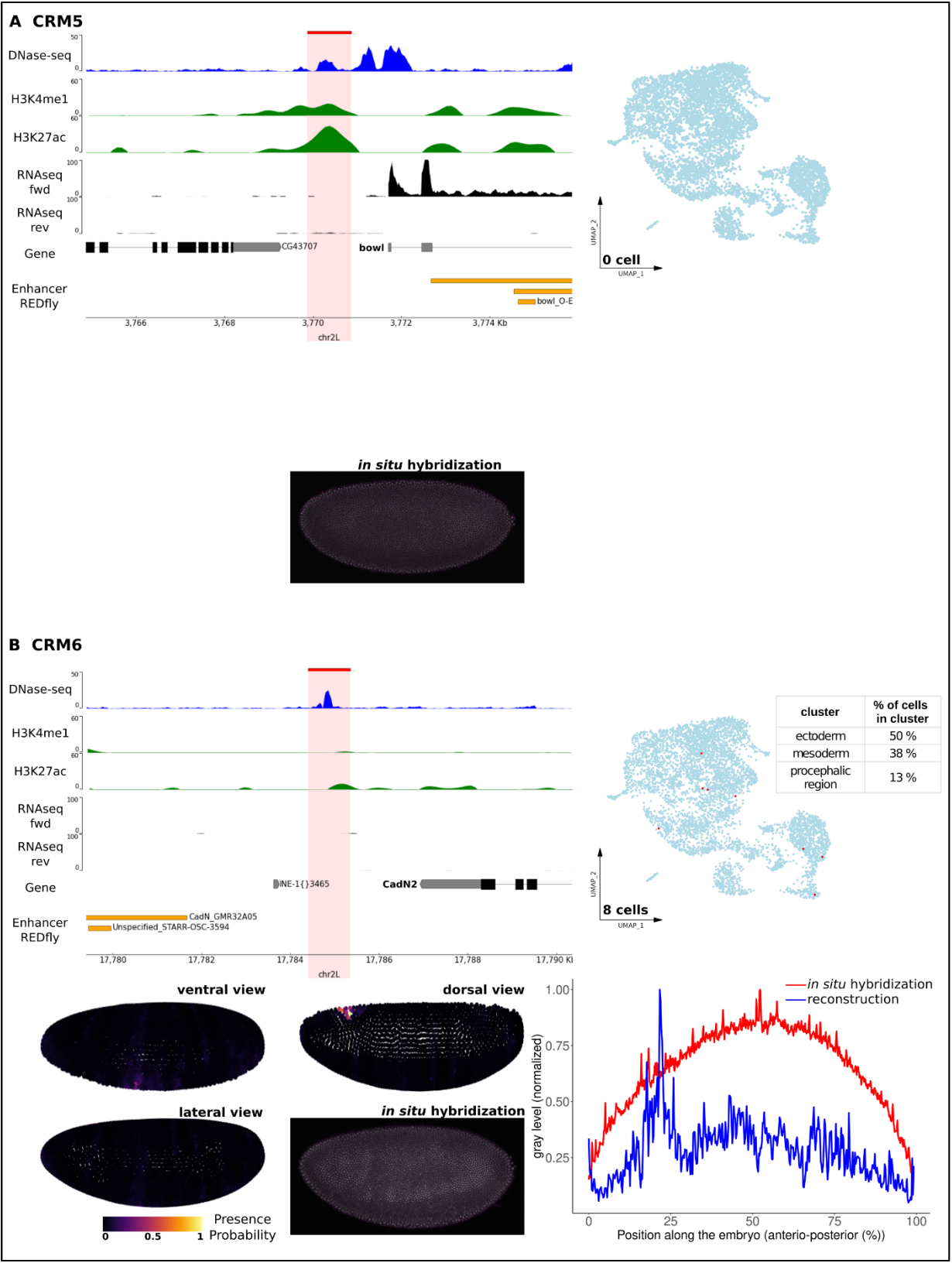

Supplementary Figure 10

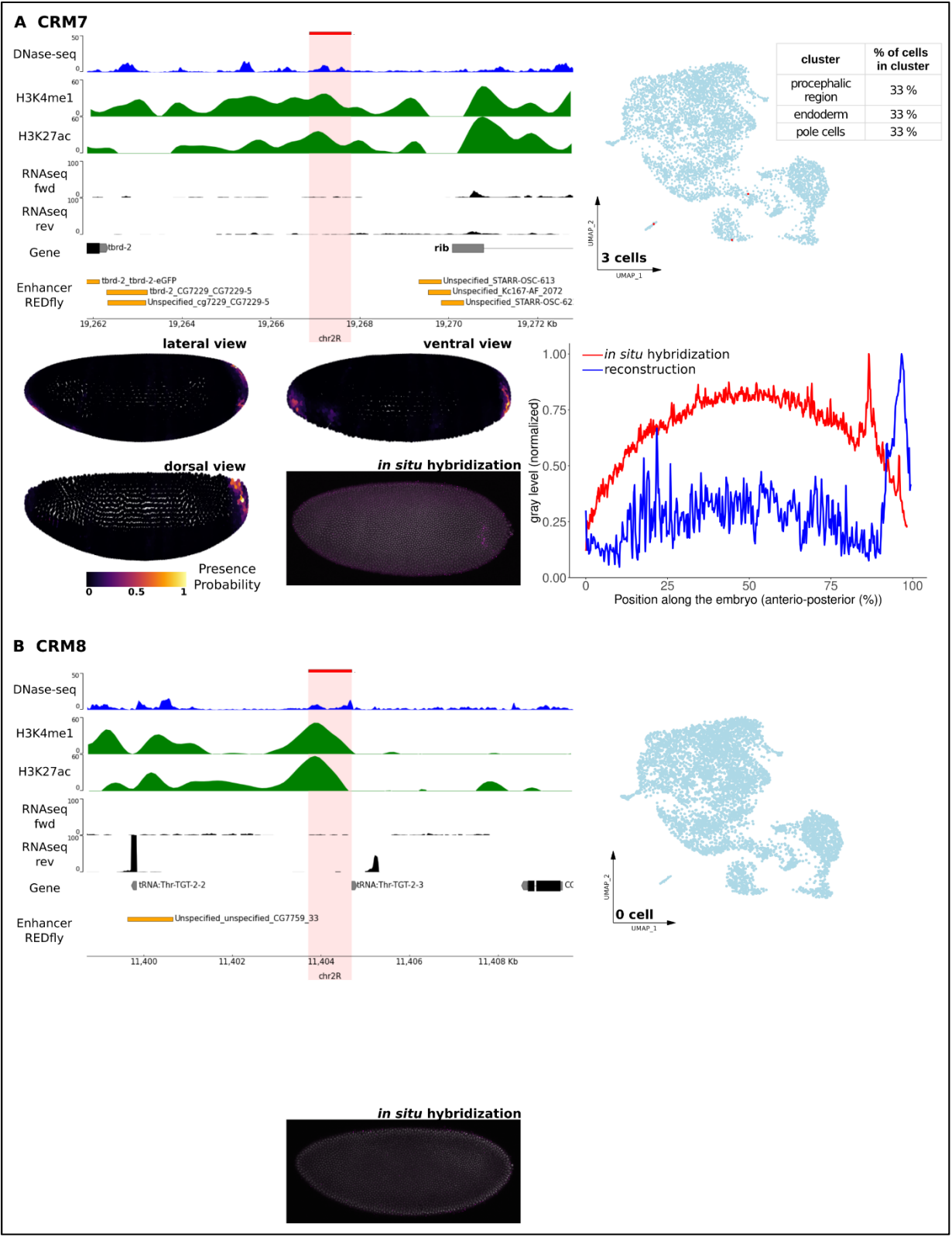

Supplementary Figure 11

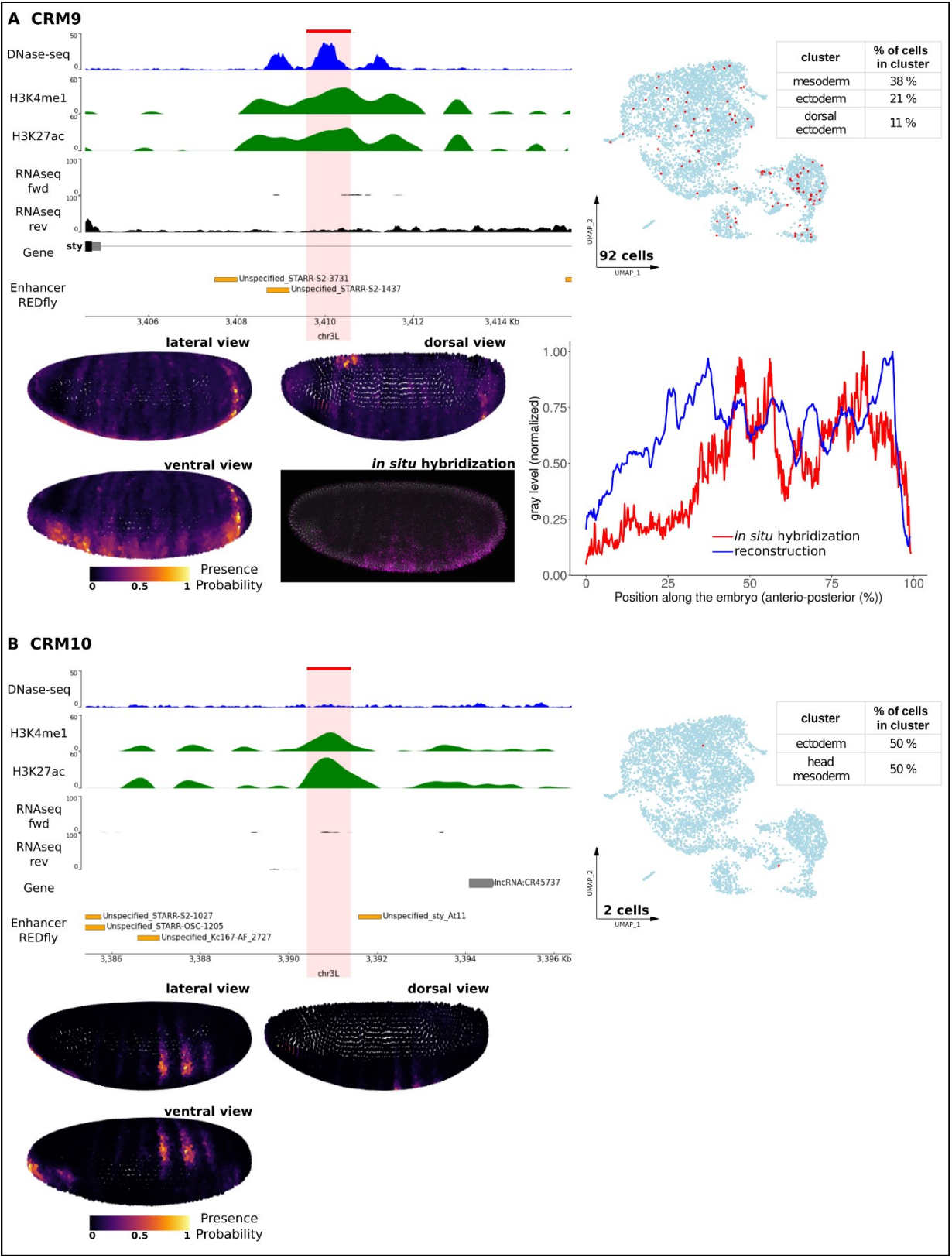

Supplementary Figure 12

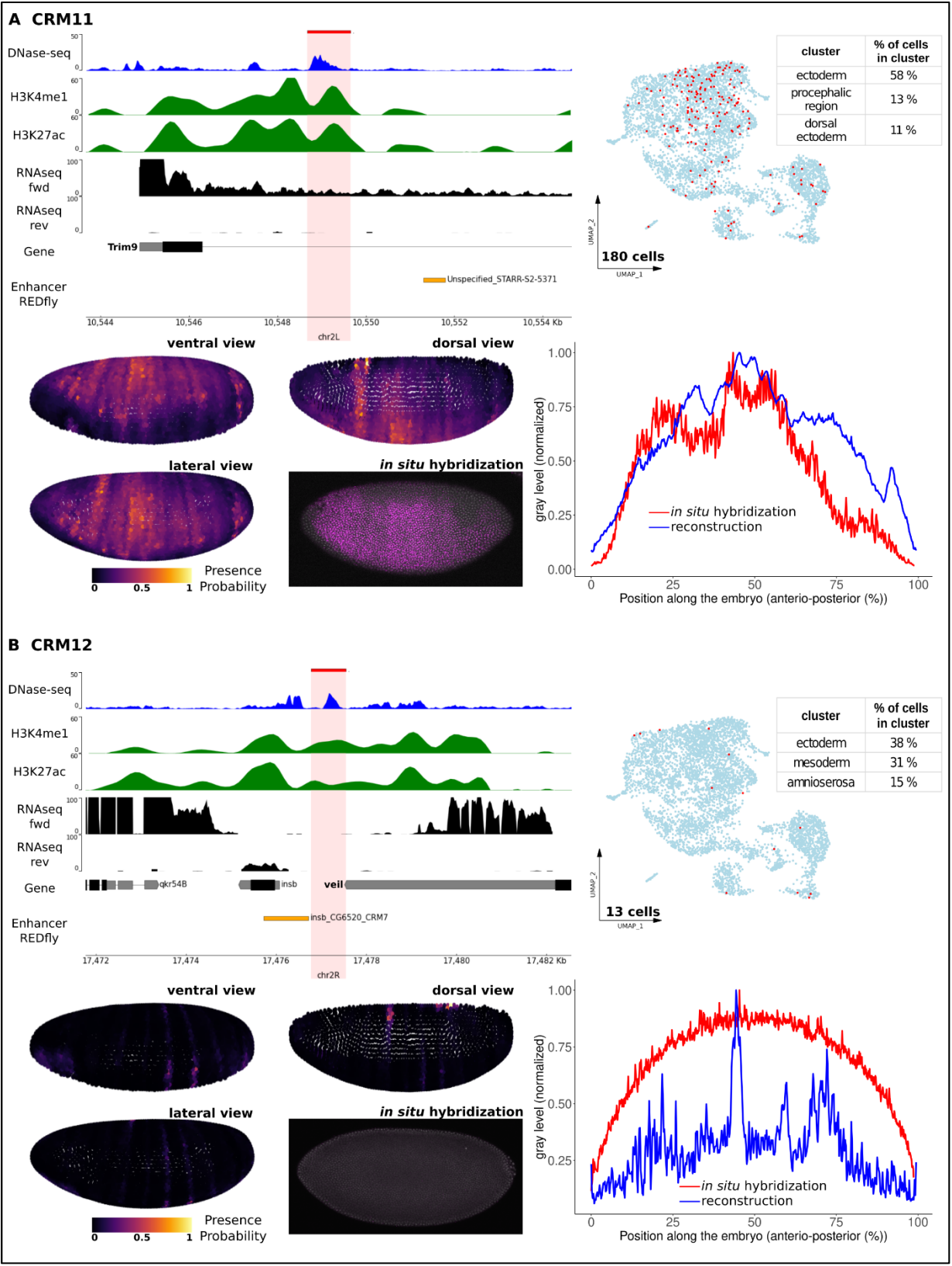

Supplementary Figure 13

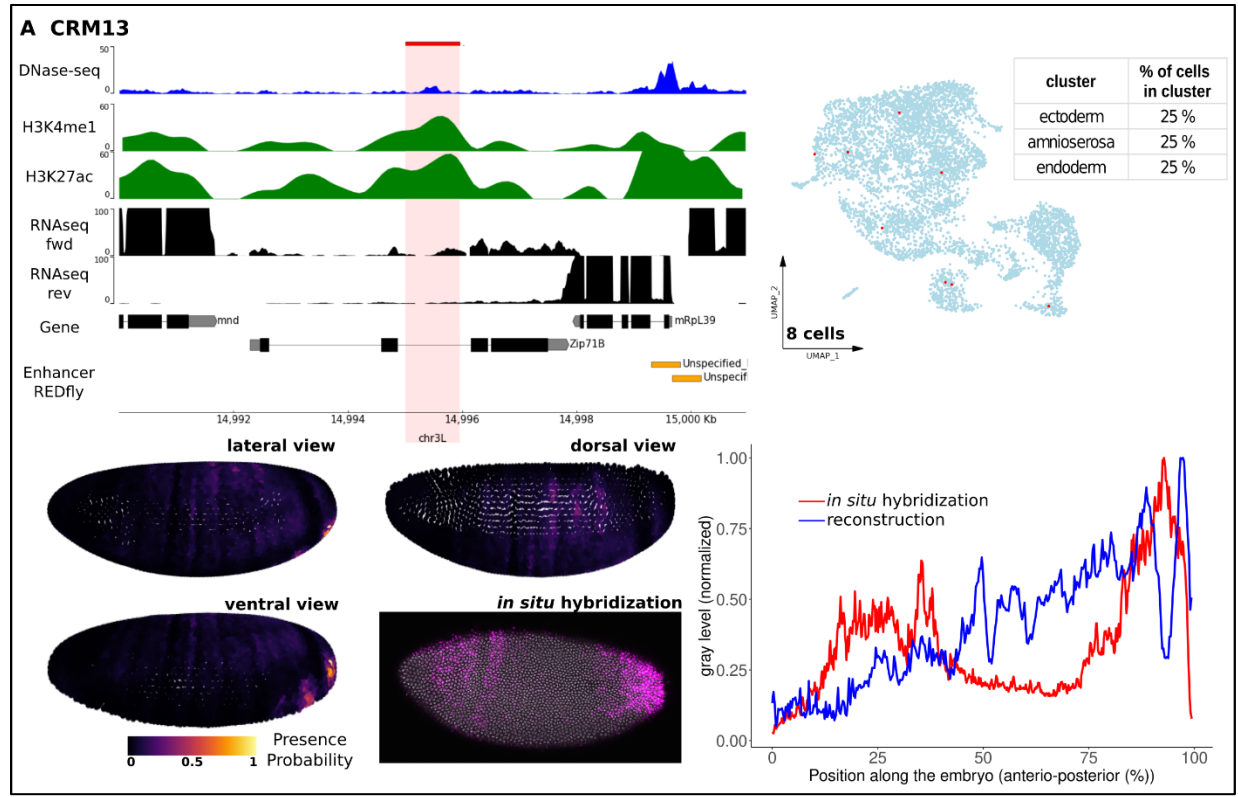

**Fig. S1-S13 Summary of all the gathered information for each candidate enhancers**

**(A-B) Top left:** Genome browser plot for each candidate region (highlighted in red). Top to bottom: DNase-seq signal at stage 5 (blue) (15); ChIP-seq signal for H3K4me1 and H3K27ac histone modifications (0-4 hours after egg lay; green) (29,30), RNA-seq signal (2-4 hours after egg lay; black) (31). The location of nearby genes and characterized enhancers from the REDfly database (orange) (32) are indicated. **Top right:** UMAP representing the candidate enhancer. Cells with an active enhancer are shown in red. The table displays the top three clusters where the enhancer is most prevalent, with percentages indicating the percentage of cells in which the enhancer is active for each cluster. **Bottom left:** reconstruction of enhancer activity obtained by plotting the probability of enhancer presence at each position in the embryos. The color code indicates the probability of presence from dark - low probability, to yellow - high probability. Three different views are shown. An *in situ* hybridization image, performed in the lab (except for vnd\_743 (46)), is included for visual comparison with the reconstruction. **Bottom right:** comparison between the profiles of average expression levels across the embryo from *in situ* hybridization experiments (red) and the reconstructions (blue).

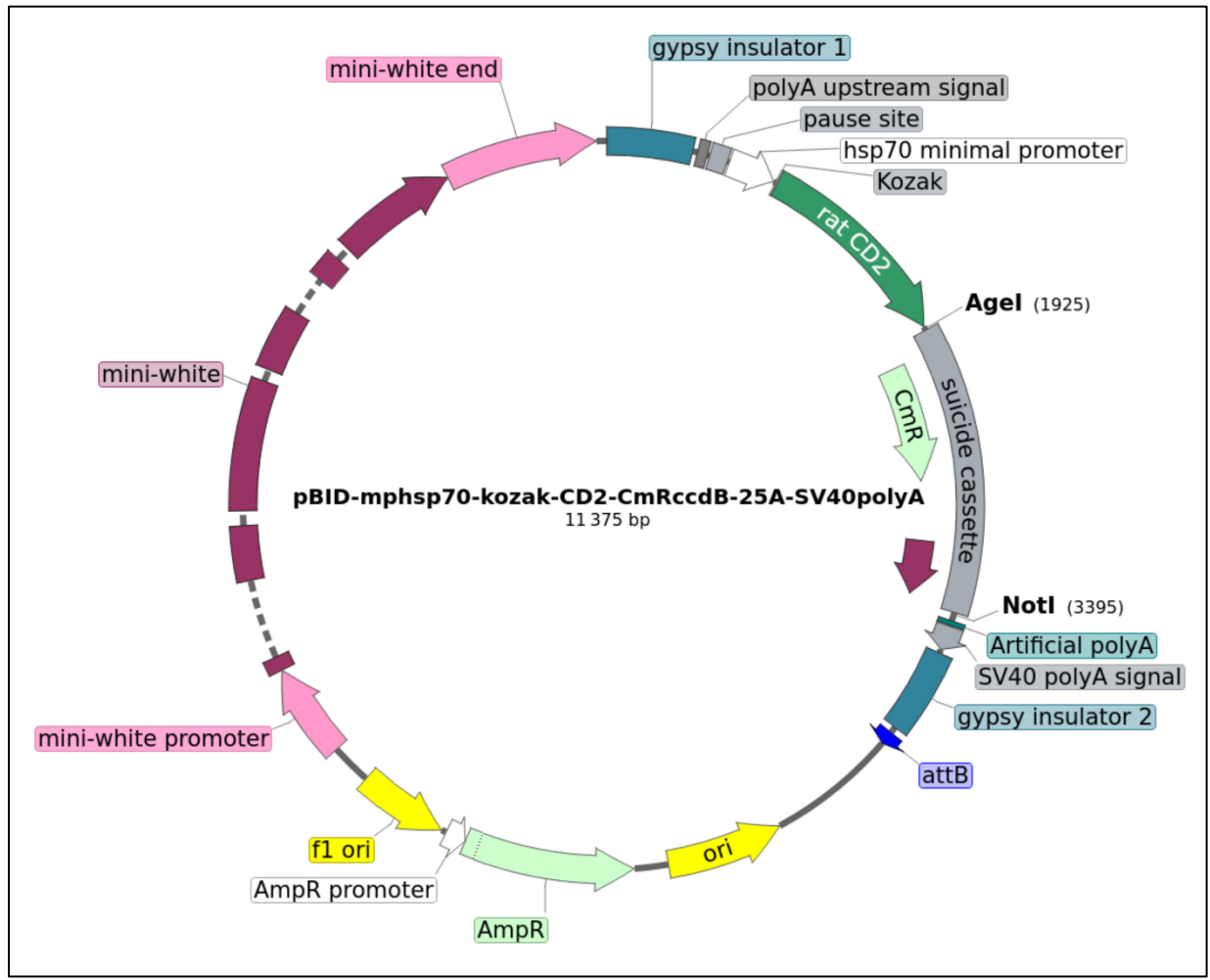

**Fig. S14 Plasmid map**

Map of the plasmid pBID-mphsp70-kozak-CD2-CmR/ccdB-25A-SV40polyA, showing key genes and selected restriction endonuclease sites. Indicated on the map are: *ori*, origin of DNA replication; *AmpR*, ampicillin resistance gene; *CmR*, chloramphenicol resistance gene; *ccdB*, bacterial toxin that poisons DNA gyrase; *mini-white* gene used for screening red-eyed flies; *rat CD2* gene used as a reporter in spatial-scERA and as target for *in situ* hybridization. Insertion in the genome is allowed by the presence of an *attB* site.

Supplementary Figure 15

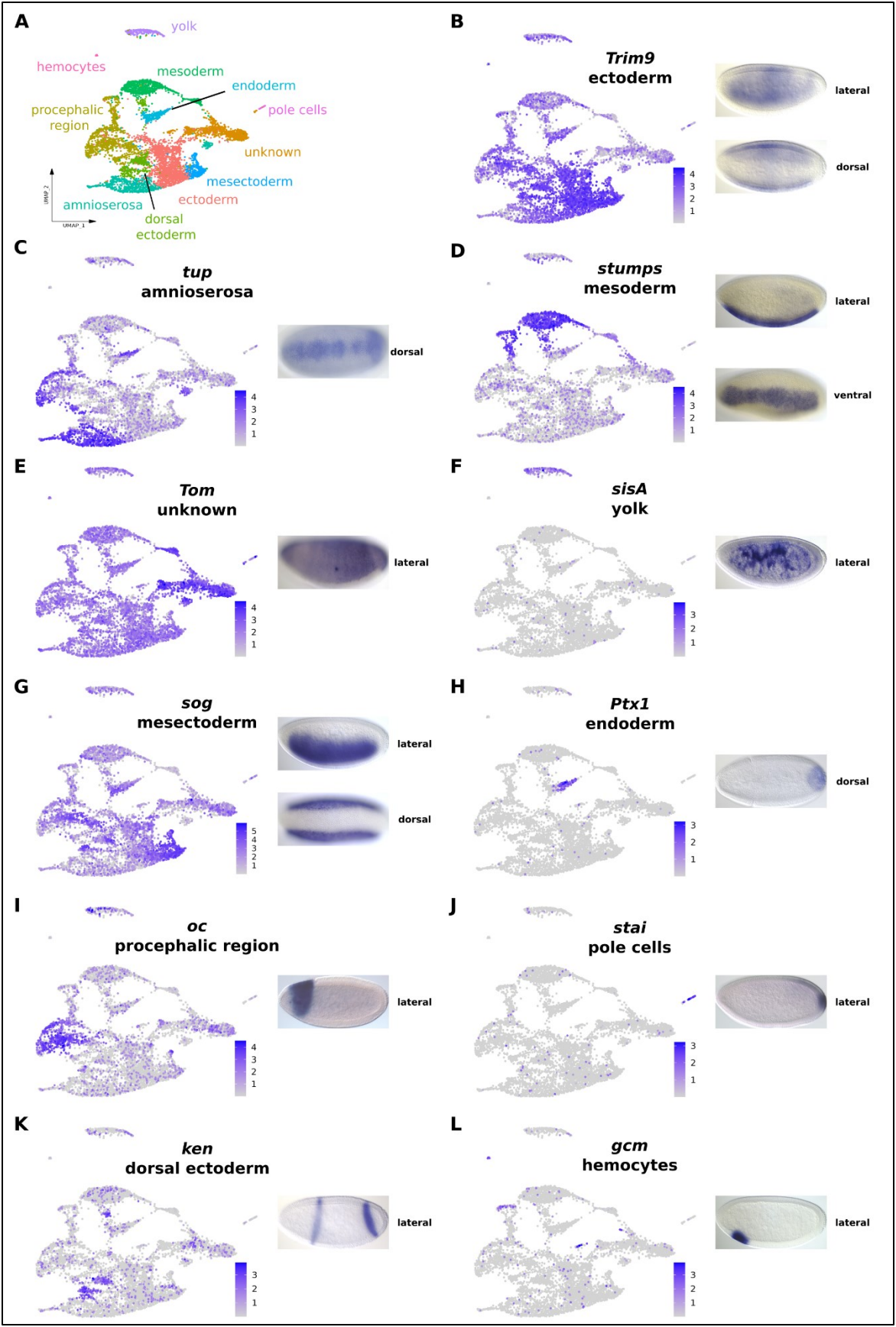

**Fig. S15 Examples of differentially expressed genes for each cluster represented on the UMAP**

(A) Uniform Manifold Approximation and Projection (UMAP) of all cells (n=6,588) from the scRNA-seq dataset, showing the 11 identified cell types. (B-L) Examples of representative genes for each of the 11 clusters. The color of the dot represents the expression level of a given gene (log2 fold change). The corresponding *in vivo* expression patterns obtained from the BDGP *in situ* hybridization database (ref) are shown on the right.

Supplementary Figure 16

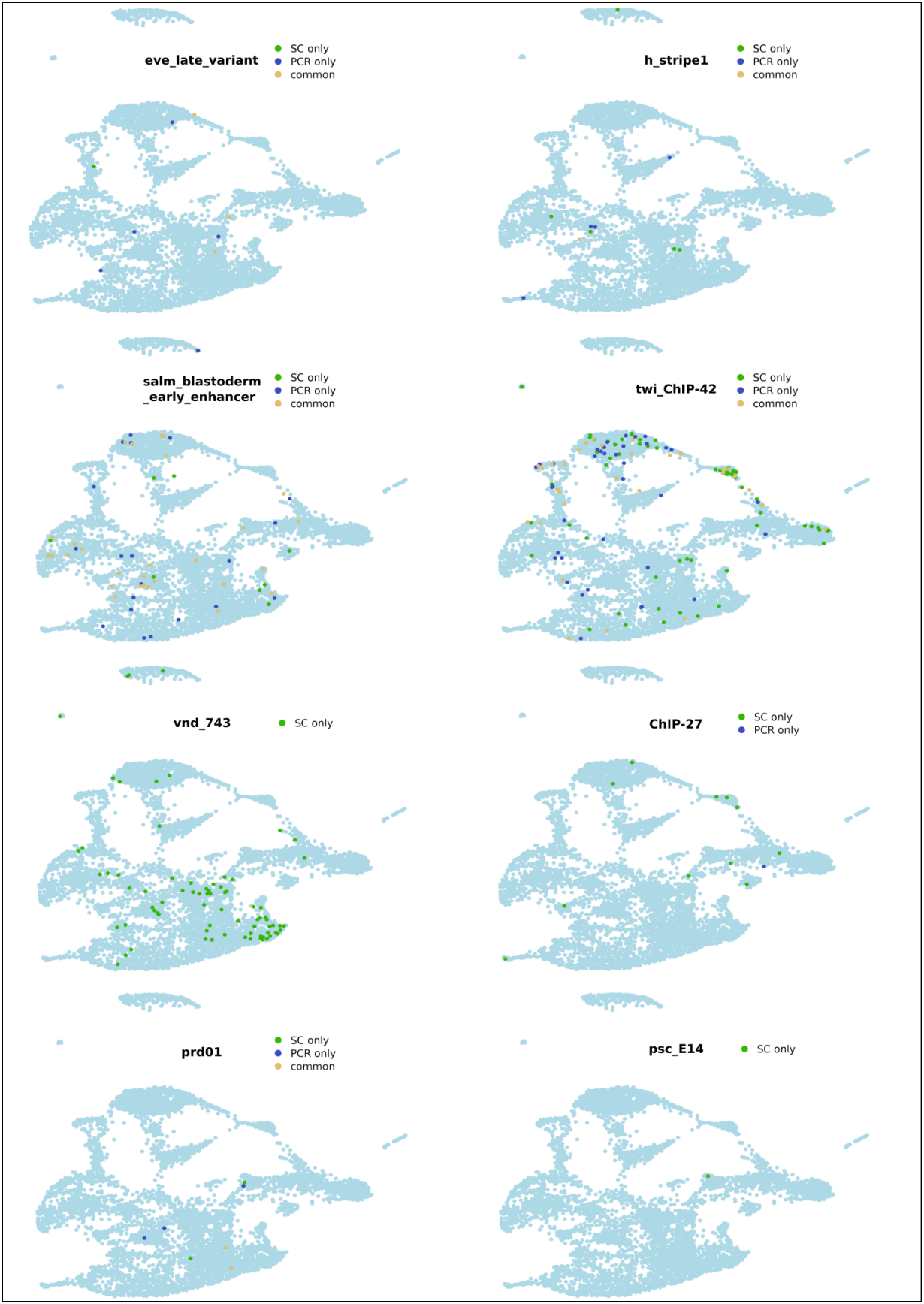

Supplementary Figure 17

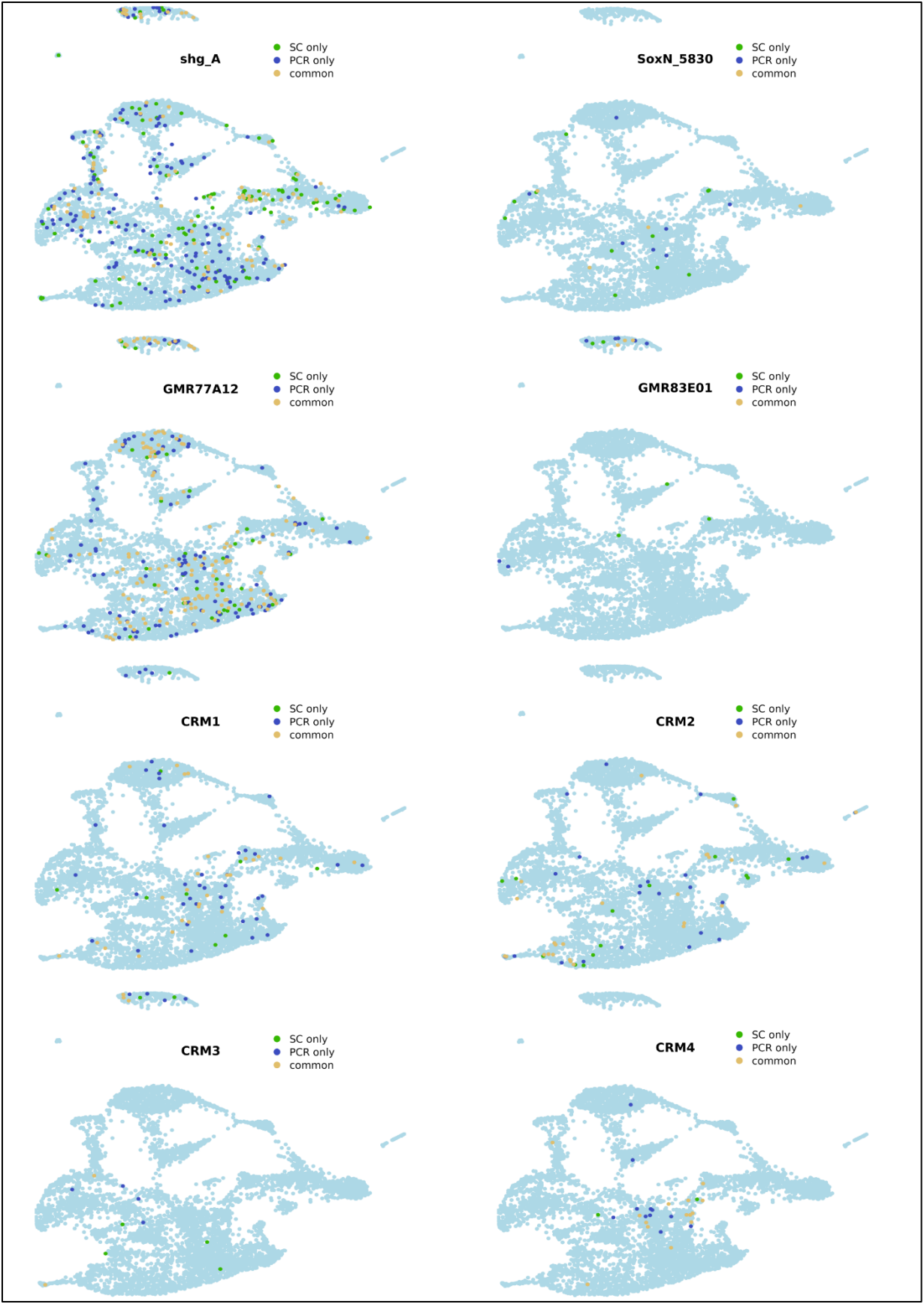

Supplementary Figure 18

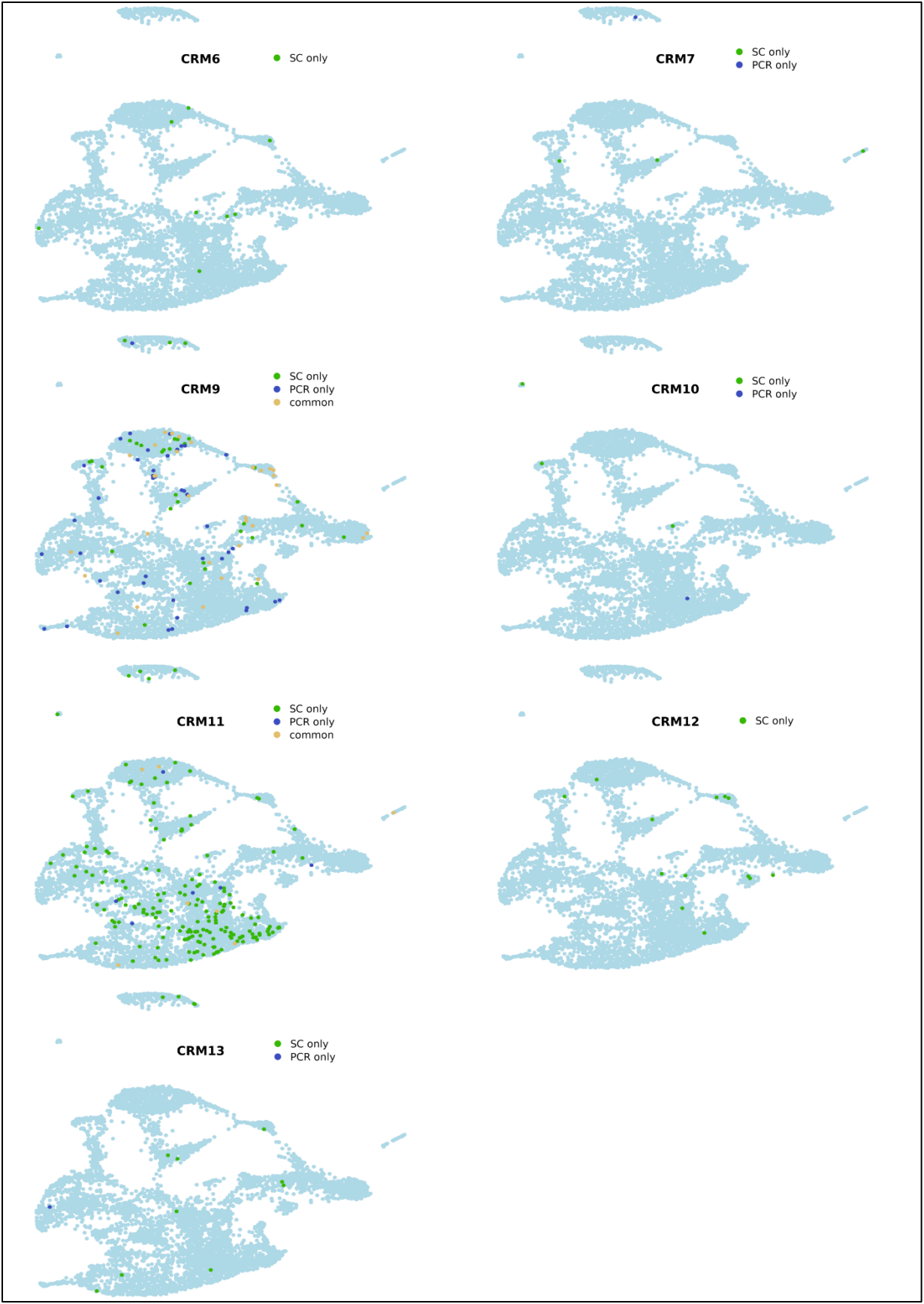

**Fig. S16-S18 Cells captured by scRNAseq and targeted PCR**

For each enhancer, the Uniform Manifold Approximation and Projection (UMAP) displays the cells discovered exclusively by scRNA-seq analysis (green), those found only by targeted PCR sequencing (blue), and those detected by both strategies (yellow).

Supplementary Figure 19

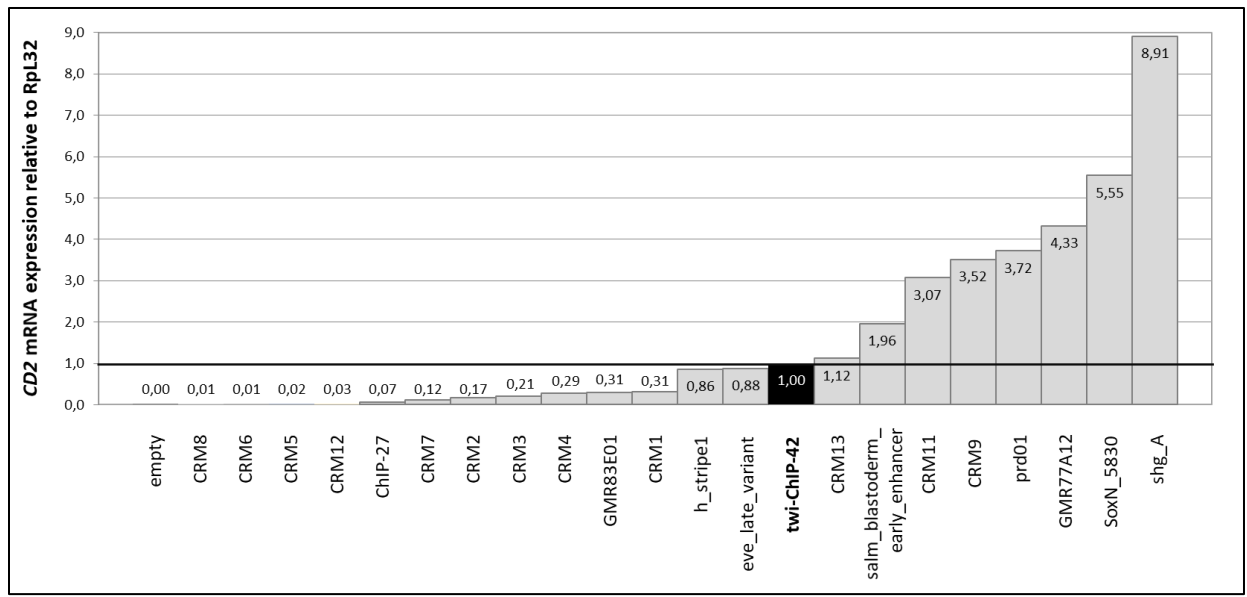

**Fig. S19 Enhancer activity in homozygous embryos assessed by RT-qPCR**

Enhancer activity was assessed in embryos carrying two copies of the same enhancer-reporter construct. The activity was quantified by measuring mRNA expression levels of the reporter gene *CD2* by RT-qPCR in embryos at 2.5 to 3.5 hours after egg lay, normalized to the expression of the housekeeping gene *RpL32*. The activity of the *twi*\_ChIP-42 enhancer (represented in black) was set to a relative expression level of 1, serving as a reference for normalizing the activities of the other enhancers.

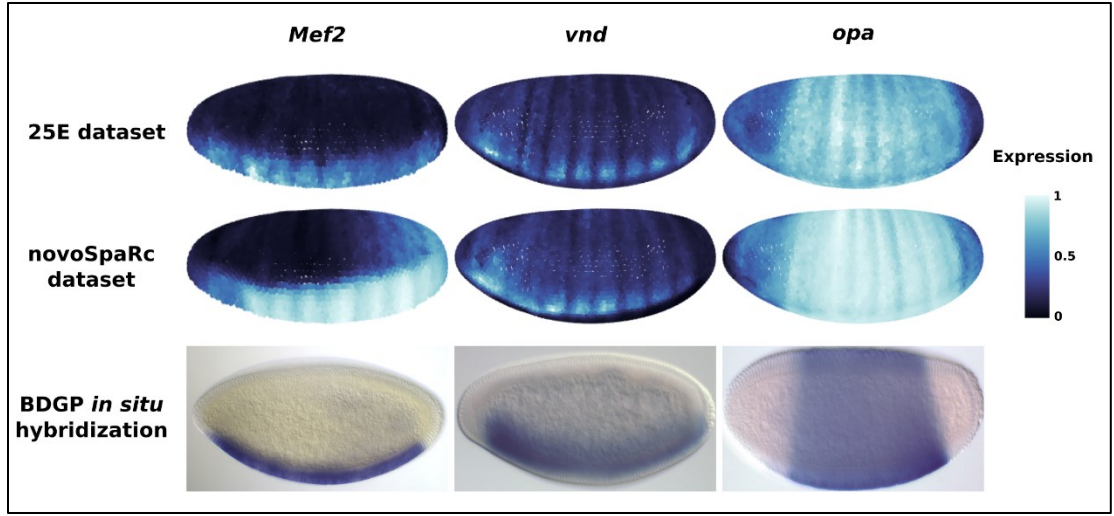

**Fig. S20 NovoSpaRc amplifies stripe patterns**

Expression patterns of three representative genes reconstructed by novoSpaRc using our dataset (top) and the scRNA-seq dataset provided with novoSpaRc (middle) (28) are compared to BDGP *in situ* hybridizations (37) (bottom). Expression levels in the reconstructions range from dark (no expression) to light blue (high expression).

Supplementary Figure 22

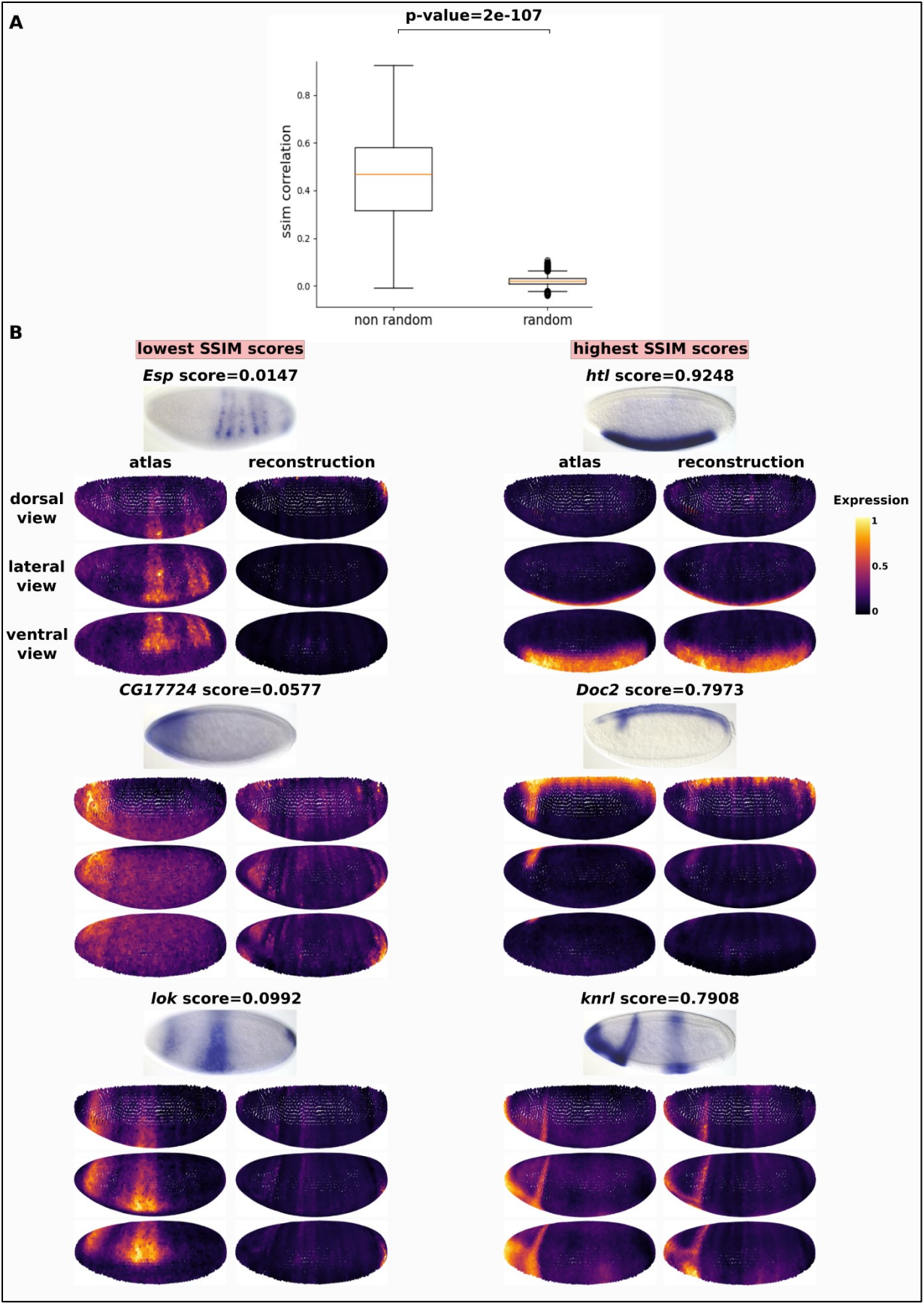

**Fig. S21 Statistical analysis of the reconstruction based on the atlas' genes**

(A) Boxplots showing the SSIM (Structural Similarity Index Measure) score for the reconstruction of all atlas genes using leave-one-out cross-validation, compared to the SSIM score of a random reconstruction model. Median is depicted as an orange bar, while the box extremities are Q1 and Q3 respectively. The p-value from the Kolmogorov-Smirnov test between the two distributions is displayed at the top of the plot.

(B) Comparison of gene expression patterns from the reconstruction with leave-one-out cross-validation against their original expression in the atlas and BDGP *in situ* hybridization data (37). The left panel shows the three genes with the lowest SSIM score, while the right panel shows the three genes with the highest score.

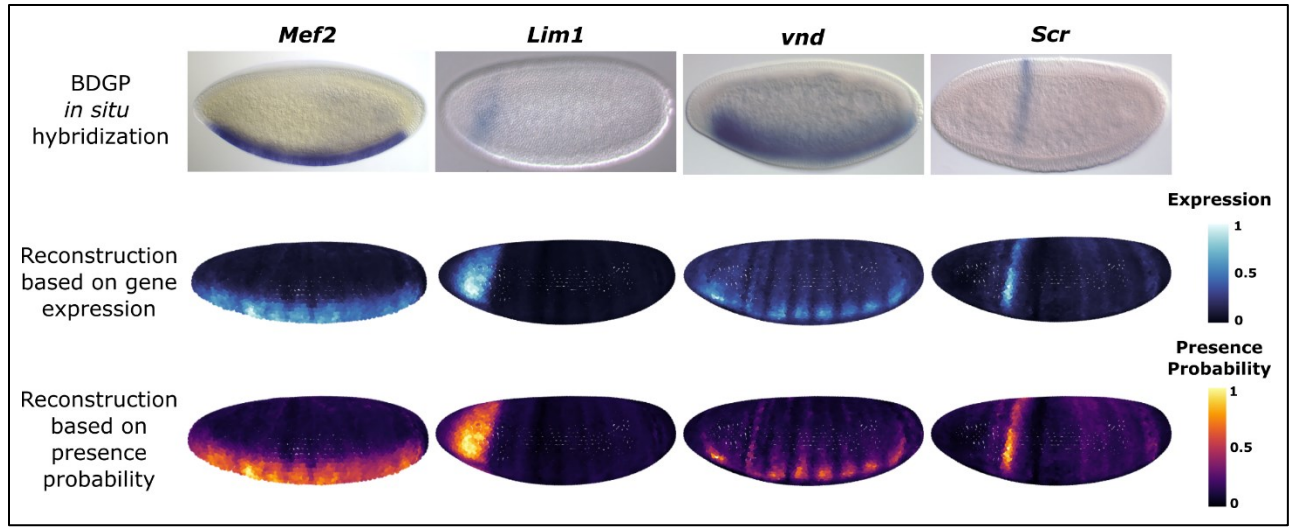

**Fig. S22 Comparison of expression patterns obtained from reconstructions based on gene expression or from predicted probability of presence**

BDGP *in situ* hybridization (top) (37) and novoSpaRc reconstructions (middle and bottom) for four different genes. The middle panels display reconstructions based on gene expression across the embryo, with dark indicating no expression and light blue indicating high expression. The bottom panels show reconstructions based on the probability of presence of each cell expressing the gene, with dark representing low probability and yellow representing high probability.

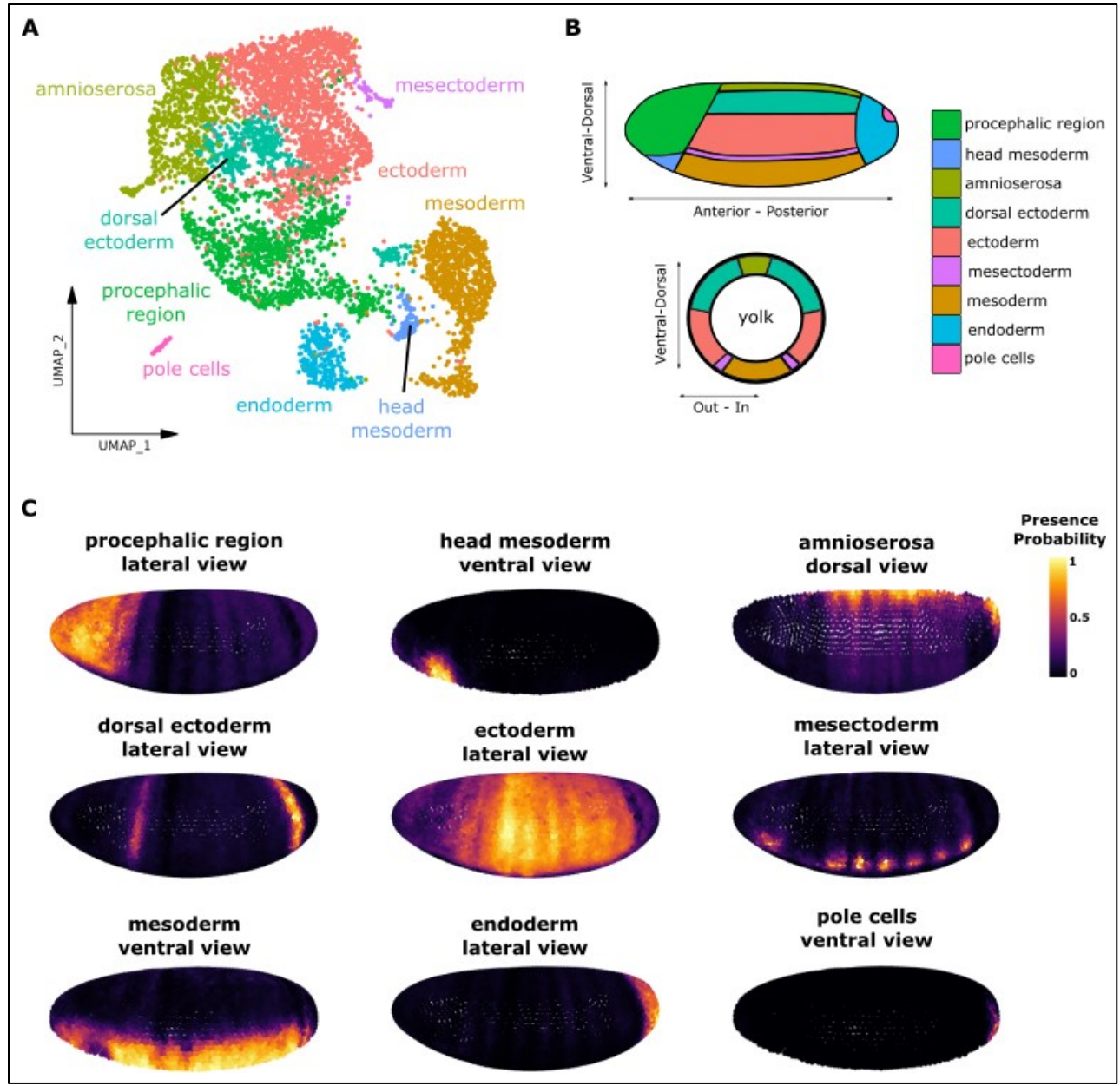

**Fig. S23 Prediction of spatial location of scRNA-seq clusters**

(A) UMAP of the reduced version (n=5,475 cells) of the scRNA-seq dataset with the 9 identified cell-types. (B) Schematic of a stage 6 *Drosophila* embryo displaying the expected location of the cell-types present in the scRNA-seq dataset. (C) NovoSpaRc reconstruction with the probability of presence of each scRNA-seq cluster.

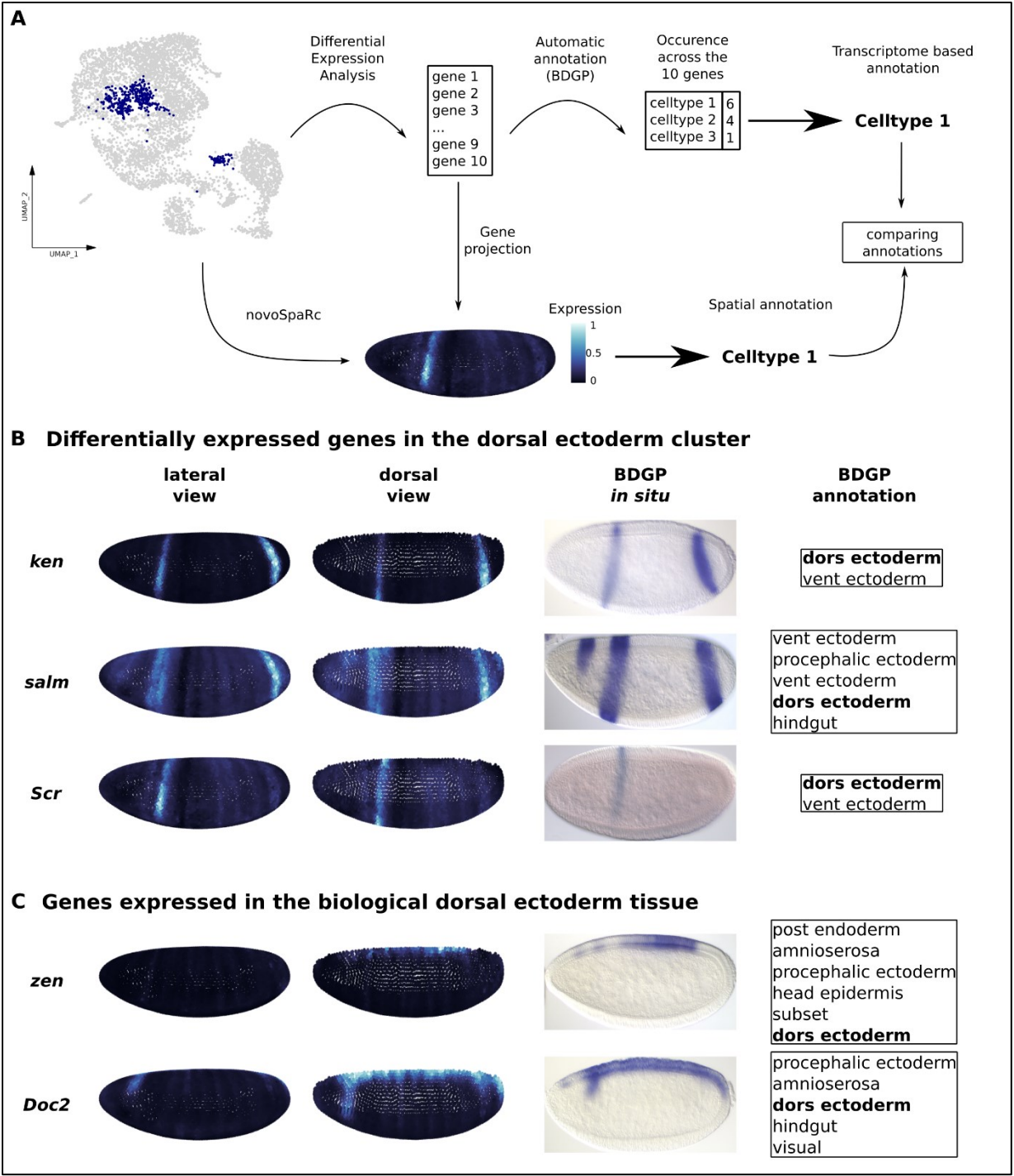

**Fig. S24 Cluster annotation strategy**

(A) Schematic of the cluster annotation strategy used during scRNA-seq analysis. For each cluster, a list of the 10 most differentially expressed genes was compiled. The most common cell type among these genes, as identified using BDGP annotations (37), was selected for cluster annotation. The reconstructed expression patterns from novoSpaRc were then used to compare and refine the scRNA-seq annotations.

### Supplementary Figure 24

**(B-C)** Examples of genes most differentially expressed in the dorsal ectoderm cluster **(B)** or known to be expressed in the dorsal ectoderm **(C)**. NovoSpaRc reconstructed expressions are shown from lateral and dorsal views. The corresponding BDGP *in situ* hybridization images and their annotations are displayed on the right

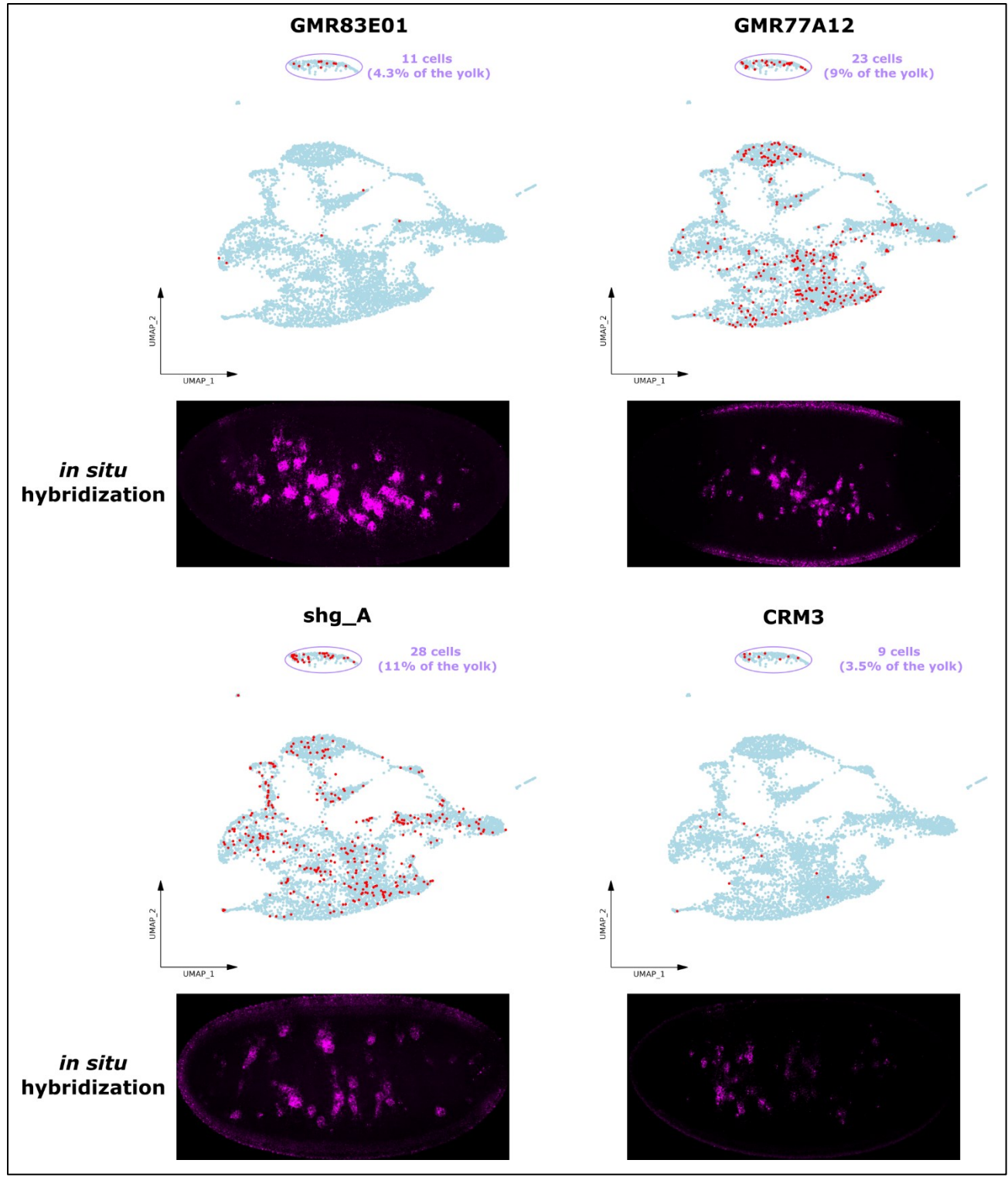

**Fig. S25 Spatial-scERA accurately captures yolk-specific enhancer activity**

(A-D) Yolk activity of four enhancers as observed in the UMAP (top) and *in situ* hybridization (bottom). The yolk cluster is circled and the number of cells carrying an active enhancer are indicated in purple

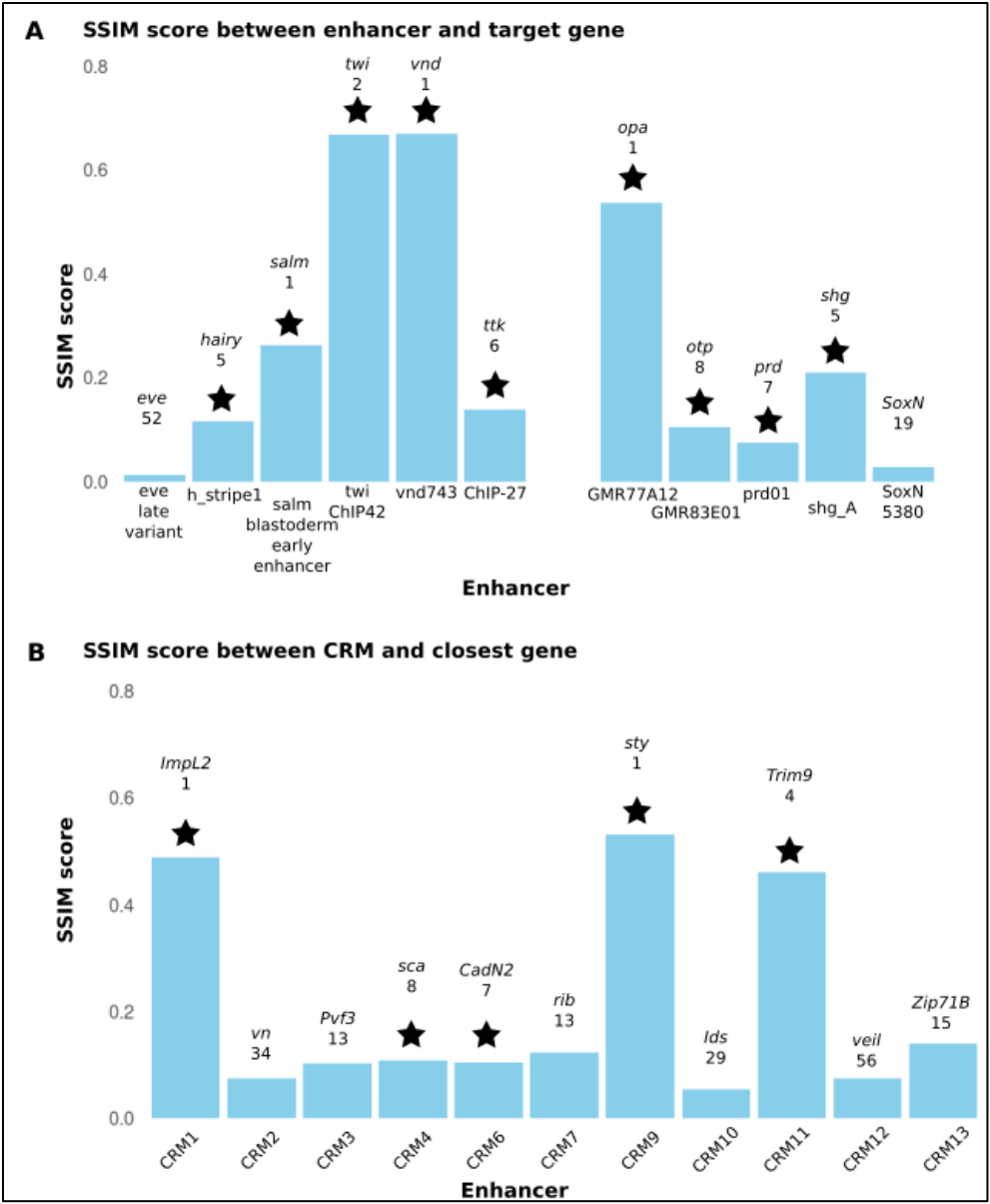

**Fig. S26 Comparing the enhancer reconstructions and the expression of the target or closest gene using the SSIM score**

**A** Histograms of the SSIM score between each previously characterized enhancer (left: at stage 6; right: at another stage) and their known target gene. **B** Histograms of the SSIM score between each of the uncharacterized enhancers (except CRM5 and CRM8 which are not active) and their closest gene (as defined in Table 1). **(A-B)** The SSIM score varies from 0 (no correlation) to 1 (full correlation). The stars indicate that the SSIM score observed for this enhancer- gene pair is among the top 10 best scores for this enhancer.

Supplementary Figure 27

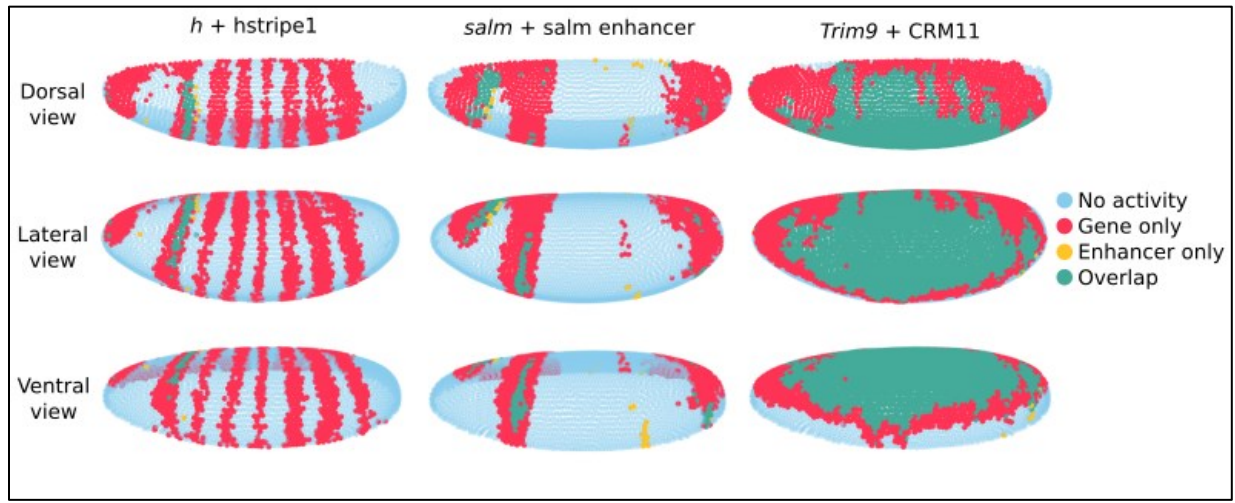

**Fig. S27 Overlap between enhancer activity and gene expression reconstructions**

Spatial reconstruction of both the enhancer (yellow) and the target or closest gene (red) for three pairs of enhancer-gene (left: *h* and *h\_stripe1*; middle: *salm* and *salm\_blastoderm\_early\_enhancer*; right: *Trim9* and CRM11). The overlap between the activity of the enhancer and the expression of the gene is shown in green.

Supplementary Figure 28

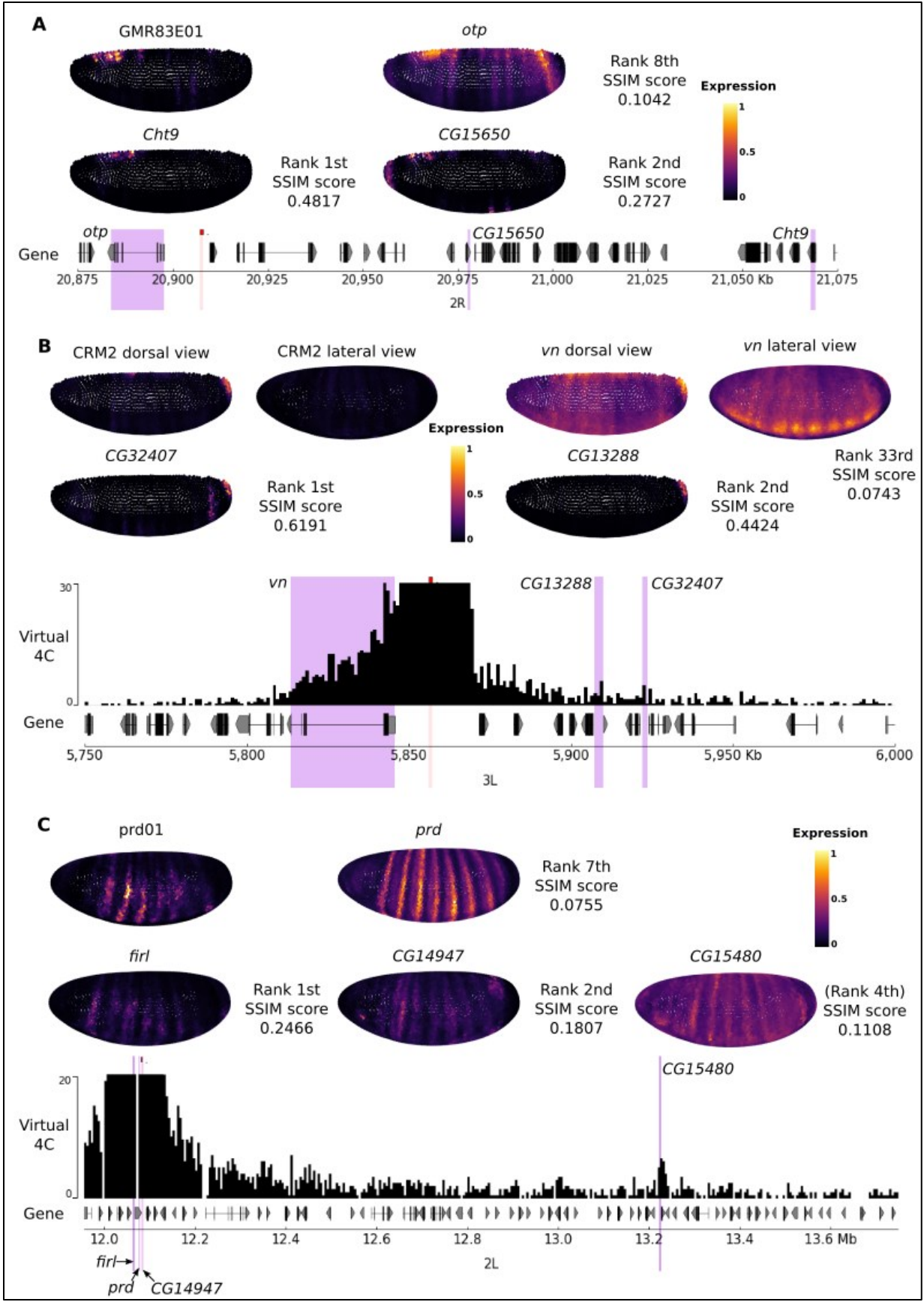

**Fig. S28 The SSIM score identifies putative new target genes regulated by the prd01, CRM2, and GMR83E01 enhancers**

**(A-C)** Spatial reconstructions of each enhancer and of several top-ranked genes based on the SSIM score (**A**: GMR83E01; **B**: CRM2; **C**: prd01). For each gene, the SSIM score and its rank among all genes within a 600kb window around the enhancer are indicated. (**B-C**) Virtual 4C maps showing interactions between CRM2 and the *CG13288* and *CG32407* genes (**B**) and between the prd01 enhancer and *CG15480* gene (**C**). Genes of interest are highlighted in purple, while enhancers are marked in red.
